# A BALB/c IGHV Reference Set, defined by haplotype analysis of long-read VDJ-C sequences from F1 (BALB/c / C57BL/6) mice

**DOI:** 10.1101/2022.02.28.482396

**Authors:** Katherine JL Jackson, Justin T Kos, William Lees, William S Gibson, Melissa Laird Smith, Ayelet Peres, Gur Yaari, Martin Corcoran, Christian E. Busse, Mats Ohlin, Corey T Watson, Andrew M Collins

**Affiliations:** Garvan Institute of Medical Research, Darlinghurst, NSW, Australia; Department of Biochemistry and Molecular Genetics, University of Louisville School of Medicine, Louisville, KY, USA; Institute of Structural and Molecular Biology, Birkbeck College, University of London, London, UK; Centre for Human-Centered Computing and Information Science, Institute for Systems and Computer Engineering, Technology and Science, Porto, Portugal; Faculty of Engineering, Bar Ilan University, Ramat Gan, Israel; Department of Microbiology, Tumor and Cell Biology, Karolinska Institutet, Stockholm, Sweden; Division of B Cell Immunology, German Cancer Research Center, Heidelberg, Germany; Department of Immunotechnology, Lund University, Lund, Sweden; School of Biotechnology and Biomolecular Sciences, The University of New South Wales, Sydney, NSW, Australia

**Author notes:** **Correspondence:** Andrew M. Collins.

**Keywords:** BALB/c, IGHV, SMRT sequencing, haplotyping, substrains

## Abstract

The immunoglobulin genes of inbred mouse strains that are commonly used in models of antibody-mediated human diseases are poorly characterized. This compromises data analysis. To infer the immunoglobulin genes of BALB/c mice, we used long-read SMRT sequencing to amplify VDJ-C sequences from F1 (BALB/c x C57BL/6) hybrid animals. Previously unreported strain variations were identified in the *Ighm* and *Ighg2b* genes, and analysis of VDJ rearrangements led to the inference of 278 germline IGHV alleles. 169 alleles are not present in the C57BL/6 genome reference sequence. To establish a set of expressed BALB/c IGHV germline gene sequences, we computationally retrieved IGHV haplotypes from the IgM dataset. Haplotyping led to the confirmation of 162 BALB/c IGHV gene sequences. A musIGHV398 pseudogene variant also appears to be present in the BALB/cByJ substrain, while a functional musIGHV398 gene is highly expressed in the BALB/cJ substrain. Only four of the BALB/c alleles were also observed in the C57BL/6 haplotype. The full set of inferred BALB/c sequences has been used to establish a BALB/c IGHV reference set, hosted at https://ogrdb.airr-community.org. We assessed whether assemblies from the Mouse Genome Project (MGP) are suitable for the determination of the genes of the IGH loci. Only 37 (43.5%) of the 85 confirmed IMGT-named BALB/c IGHV and 33 (42.9%) of the 77 confirmed non-IMGT IGHV were found in a search of the MGP BALB/cJ genome assembly. This suggests that Adaptive Immune Receptor Repertoire sequencing (AIRR-Seq) data, but not currently-available genome assemblies, are suited to the documentation of germline IGHV genes.

## Introduction

Our understanding of B cells and the antibody response have long been informed by studies of immunoglobulin genes. For example, early genetic studies revealed processes such as somatic point mutation and affinity maturation (1, 2). Other studies focused upon the expression of particular immunoglobulin genes, in an effort to better understand the aberrant immune responses that are seen in allergic (3) and autoimmune diseases (4). Today, new insights are coming from the study of immunoglobulin gene repertoires using high-throughput sequencing.

Immunoglobulin genes are found within the genome as multiple sets of highly similar genes. The immunoglobulin heavy chain (IGH) that is the focus of this study is encoded by Variable (IGHV), Diversity (IGHD) and Joining (IGHJ) genes. Multiple genes of each type are found together within the IGH gene locus. During early B cell development, genetic recombination joins one of each of these gene types to form a functional VDJ gene (5). The VDJ gene is expressed in association with a single Constant (C) region gene to produce the immunoglobulin heavy chain, while the light chain is produced by similar processes acting on separate sets of light chain genes.

The study of human VDJ genes was facilitated first by the documentation of the complete set of germline immunoglobulin genes that are available for recombination (6, 7), and later by the documentation of allelic variation and structural variation within the human population (8–10). With this knowledge to hand, analysis of the expressed antibody repertoire became possible. It is now known that the diversity of the antibody repertoire is an outcome of processes that are stochastic, but that are nevertheless influenced by germline variants (11–13).

Adaptive immune receptor repertoire sequencing (AIRR-seq) studies now often report the sequencing of thousands and even millions of different rearranged immunoglobulin genes from a single individual. Over the last decade, such sequencing studies have transformed our understanding of the nature of the human antibody repertoire, and of fundamental aspects of the antibody response in health and disease (14–18). But despite the importance of animal models for our understanding of antibody-mediated immunity, autoimmunity and allergic disease, there have been relatively few reports of the antibody repertoires of laboratory mice. In part, this is a consequence of our lack of understanding of the germline immunoglobulin genes of the mouse, for only the genes of the C57BL/6 mouse have been comprehensively documented. Without an understanding of their germline genes, reliable analysis of the repertoires of other mouse strains is impossible.

The germline genes of the BALB/c mouse have previously been explored using AIRR-seq data and a process of gene inference (19). In such analyses, the presence of multiple examples of identical sequences within the set of VDJ gene rearrangements is used to identify each germline gene. The reliable inference of germline IGHV genes within datasets of VDJ gene rearrangements, particularly those of naïve B cells, is a trivial exercise for highly expressed genes, but it is challenging for rarely expressed genes. Many BALB/c genes appear to consistently rearrange at frequencies as low as 0.01%. Given the sequencing depth of the study by Collins and colleagues (19), many of their BALB/c germline gene inferences were only supported by a handful of rearranged VDJ gene sequences. It is therefore important that these inferences be further tested using validated inference tools (20, 21).

High-throughput AIRR-Seq using long read single-molecule real-time (SRMT; Pacific Biosciences) sequencing can cover the entire V(D)J region and extend into the CH3 exon of the constant region. It was recently applied to the study of the mouse VDJ repertoire (22). Here, we used this approach to amplify VDJ rearrangements in association with constant region genes (VDJ-C). The F1 IGHV genotype was defined from rearranged VDJ-C amplicons and IGHJ1-based haplotyping was used to explore genetic variation within the *Ighm* and *Ighg*-encoding constant region genes. Strain-defining single nucleotide polymorphisms (SNPs), including variants that are not present in the IMGT database, were identified, allowing the RAbHIT haplotyping tool (23) to be used to assign each IGHV to one or other of the parental strains. This analysis largely confirms the reported germline IGHV sets for the C57BL/6 and BALB/c strains. Only four IGHV sequences are shared by the two strains.

It is now known from studies of wild-derived and classical inbred mice that the IG gene loci are highly divergent between strains (19, 22). The divergence is so great that it is impossible to assign new mouse sequences as allelic variants of genes that have been defined by the C57BL/6 genome. Strain-specific IGHV Reference Sets should allow accurate analysis of the immunoglobulin repertoire, without reference to C57BL/6 genes, but there is presently a lack of strain-specific IGHV Reference Sets for non-C57BL/6 strains. To begin to address the need for such sets, the BALB/c sequences identified in this study were used to establish a curated BALB/c IGHV Reference Set at the Open Germline Receptor Database (OGRDB) website (https://ogrdb.airr-community.org/) (24). The errors that can arise from the use of incomplete Reference Sets in AIRR-Seq analysis are highlighted here, and we suggest that it will be critical for the IGHV repertoires of additional inbred strains to be properly documented if we are to better understand the roles of antibodies in mouse models of human disease.

## Materials and Methods

### Mice and Library Construction

RNA was extracted from BALB/cByJ x C57BL/6J F1 hybrid dissected spleens (n = 4) preserved in RNA later (Thermofisher, Cat. No. AM7020; Waltham, MA, USA). Briefly, total RNA was extracted from 30 mg of preserved spleen using the RNeasy Mini kit (Qiagen, Cat. No. 74104; Germantown, MD, USA). First-strand cDNA was generated from 1 μg of RNA for each sample using the SMARTer RACE 5’/3’ Kit (Takara Bio, Cat. No. 634858; Mountain View, CA, USA). For each sample, long-read single molecule real-time (SMRT) sequencing (Pacific Biosciences) libraries were prepared using IgM and IgG constant region primers that amplified off the first-strand cDNA synthesis product. Rearranged VDJ IGM and IGG amplicons were generated using a universal forward primer (Takara Bio;10 μM), and either an IgM-CH3 (5’-CAGATCCCTGTGAGTCACAGTACAC-3’; 10 μM) or IgG-CH3 (5’-ATGAAGTAAGAACCATCAGAGTCCAGGAC-3; 10 μM’) reverse primer. First-strand cDNA was amplified using Thermo Fisher Phusion HF Buffer (Thermo Fisher, Cat. No. F530S; Waltham, MA, USA) for 30 PCR cycles. Final amplicons were used to construct SMRTbell libraries and sequenced on either the RSII or Sequel system (Pacific Biosciences; Menlo Park, CA, USA).

### Definition of Germline Gene Reference Sets

Genotype inference tools such as TIgGER (21, 25) are able to identify previously unreported germline IGHV in AIRR-seq datasets, but they work best with reference datasets that correspond fairly closely to the genotypes under investigation. If an unreported sequence is substantially different to any sequence in the reference dataset, it may not be detected. We therefore sought to confirm the results of an earlier study (19), rather than determining the F1 genotype *de novo*. We compiled a reference set of all BALB/c and C57BL/6 genes identified in the study of Collins and colleagues, as well as all other reportedly functional C57BL/6 or BALB/c IGHV genes in the Reference Directory of the ImMunoGeneTics (IMGT) group. Some IGHV gene sequences and names in the IMGT Reference Directory have changed since the publication of the Collins study. Sequence changes mostly involve the 3’ terminal nucleotides, and are largely undocumented at the IMGT website, but were identified through archived webpages. Name changes and sequence changes are documented in **Table I**.

**Table I:**
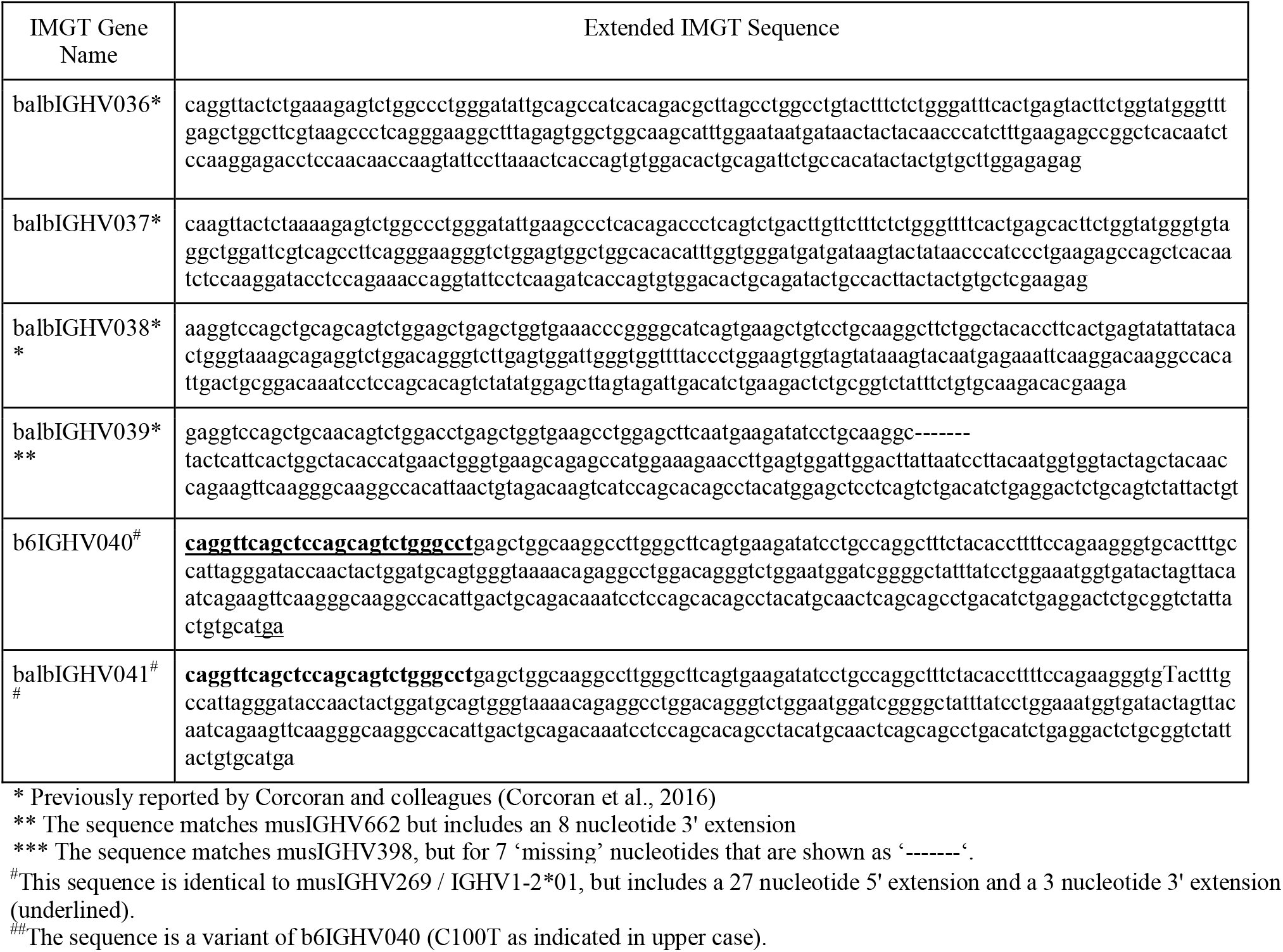
New BALB/c IGHV sequences identified in this study

The performance of annotation tools as well as germline gene inference tools can be compromised by the presence of truncated sequences in a reference set. A number of truncated genes from IMGT and other sources are annotated as being present in the BALB/c strain. To maximize the likelihood of their detection by genotyping tools, reference set sequences that appeared to have short truncations were manually extended by reference to their most similar full-length IGHV sequences (**Supplementary Table I**). Substantial extensions were required for other IGHV sequences that are referred to, by IMGT, as being ‘unmapped’. These sequences can be identified by IMGT’s ‘S’ nomenclature: for example, IGHV1S22*01. These truncations involve both 3’ and 5’ nucleotides, with as many as 57 nucleotides being missing from individual sequences (**Supplementary Table I**).

Extensions of eight of the ten ‘unmapped’ sequences were made with reference to other Genbank sequences that were exact matches to the truncated sequences (**Supplementary Table I**). In these eight cases, the Genbank sequences were recorded as being derived from BALB/c animals, or from animals of the BALB/c-derived D-limited strain (26). A ninth sequence, IGHV9S8*01, was originally reported in BALB.K mice. It was extended using the closely matching genomic sequence (AY169679) reported from the DBA strain (27). Both the BALB.K and DBA stains are closely related to the BALB/c strain. An exact match to the final ‘unmapped’ sequence, IGHV1S75*01, was found in the BALB/c genome sequence of the Sanger Mouse Genome Project (MGP) at Ensembl 104 (28). All the extended sequences were used in the knowledge that errors in the extensions could be readily identified in this study as part of genotyping and the search for novel alleles using inference processes.

The incorporation of all these changes led to the definition of a reference set made up of modified and unmodified IMGT and other sequences called the Combined and Extended (ComEx) Reference Directory. It was used as the starting point for this study. Over the course of the study, the ComEx Reference Directory was modified by the inclusion of additional inferred novel alleles, and final analysis was based upon this modified version of the Reference Directory.

### Processing of long read VDJ-C datasets

UMIs were extracted from PacBio SMRT-seq Q30 fastq files and primers were trimmed using MaskPrimer from Immcantation’s presto package (v0.5.4) (29). Reads were aligned using IgBLAST (v1.14) (Ye et al., 2004) to the ComEx IGHV and IMGT’s mouse IGHD and IGHJ reference directory (reference obtained from IMGT 16-01-2020) to generate AIRR-formatted output (-outfmt 19). Change-O databases were generated using the MakeDB command from the Change-O tool (v 1.1.0) with --failed option to retain both the ‘pass’ and ‘fail’ subsets (30).

The *Ighc* portion of each read was extracted based on the position of the final IGHJ nucleotide in the IgBLAST output, with the *Ighc* part running from 1 nucleotide downstream from the IGHJ through to the end of the primer trimmed read. Upon definition of the strain-specific spliced constant region exons for the strains, the *Ighc* portions were insertion/deletion (indel) corrected via blastn alignment (31) and processing with a custom script. This permitted positional extraction of the *Ighm* and *Ighg2b* SNPs.

### Defining the F1 Genotype

Change-O databases for the IgM datasets, both pass and fail, were merged with the extracted *Ighc* sequences and combined to generate a single F1 dataset. VDJ-C reads that were 5’ truncated or which lacked a CDR3 or an IGHJ gene call were removed, and the dataset was collapsed to unique VDJ nucleotide sequences. TIgGER’s FindNovelAlleles was called on this F1 IgM dataset with germline_min = 50 and min_seqs = 10.

Before proceeding to genotyping, novel alleles were added to ComEx. IgBLAST was re-run and new Change-O databases were created. Genotyping of the IgM F1 datasets was performed using TIgGER (v1.0.0) (21) with the inferGenotypeBayesian function. Following genotyping, alleles were reassigned using the reassignAlleles function as input for haplotyping. The F1 genotype and associated fasta set were exported from TIgGER and processed with OGRDBstats (https://github.com/airr-community/ogrdbstats), providing data for the clarification of the 3’ ends of the genes.

### Defining strain specific genes via haplotyping

The BALB/c strain carries the IGHJ1*01 allele, which differs from the C57BL/6 IGHJ1*03 allele at a single SNP (32, 33). IGHJ1-based haplotyping was performed to explore variability between strains in the constant region exons. Reads were filtered for IGHJ1 usage and then subsets were created based on either IGHJ1*01 or IGHJ1*03 usage. Unique *Ighc* sequences associated with the *01 and *03 VDJs were sorted according to their frequency. The top *Ighm* sequence associated with each IGHJ1 allele was selected and aligned to the C57BL/6 and BALB/c genomes at ensembl (ensembl.org). The IGHJ1-based haplotyping was repeated on the IgG F1 dataset, but with the top 2 sequences for each IGHJ1 allele extracted to account for IgG2a and IgG2b in BALB/c and IgG2b and IgG2c in C57BL/6.

Identification of strain-defining SNPs in the *Ighm* gene allowed haplotyping of the IGHV loci of the two strains to be determined using unmutated IgM-associated VDJ sequences. Haplotyping was performed using RAbHIT (v0.1.8) (23, 34). Input to haplotyping was the TIgGER reassigned F1 IgM dataset joined to the IgM SNP-typed data in the form of a c_call with format IgM*B6 or IgM*BALB. Sequences lacking an *Ighc* call were excluded, as were those with ambiguous calls (for example, the same VDJ nucleotide sequence that was associated with both IgM*B6 and IgM*BALB). Haplotyping was also limited to unmutated, unique sequences with read counts greater than 1. The RAbHIT function createFullHaplotype was called with the following parameters: toHap_col = v_call, hapBy_col = c_call, hapBy = IgM, kThreDel = 0.1, relative_freq_priors = FALSE and single_gene = TRUE. IGHV8 haplotypes were plotted with the plotHaplotype function using a custom genes_order and removeIGH = FALSE.

C57BL/6 and BALB/c IGHV sets were output from RAbHIT haplotypes using the TIgGER germline inferred database. Prior to incorporation into the OGRDB BALB/c Reference Set, gene ends were manually reviewed using analysis of terminal nucleotide triplets.

To assess the consequences of the use of an incomplete Reference Set for the analysis of BALB/c data, we analyzed publicly available datasets from the study of Corcoran and colleagues (20). The data was generated from peripheral blood lymphocytes using 5’ RACE and Illumina MiSeq sequencing. The animals were of the BALB/cJ substrain, giving an opportunity to also explore IGHV gene and repertoire differences between the BALB/cByJ and BALB/cJ substrains. IgBLAST was first run using the IMGT Reference Set and then using the OGRDB BALB/c Reference Set, noting the number of mismatches in each IGHV alignment.

## Results

### Long read VDJ-C sequencing of F1 mice

IgM CH3 primed libraries were sequenced from four F1 mice (J1, J2, J3, J4), with replicate sequencing from mouse J2 (three replicates) for a total of 6 IgM datasets. IgG CH3 primed libraries were sequenced from the same four animals. IgG from J1 generated a low read count and was excluded from analysis.

A combined total of 383,550 IgM reads and 67,237 IgG reads passed primer trimming. Complete VDJ-C amplicons were found in 117,262 IgM reads and 59,217 IgG reads. Filtering of the IgM dataset retained 51,988 unique, full length IgM sequences that were used for novel allele predictions and a subset of 36,294 unmutated sequences were used for genotyping of the F1. Haplotyping utilized 7,896 sequences with highly confident *Ighm* constant region SNP calls.

### Novel alleles identified in F1 dataset

The analysis of 51,988 reads aligned against the ComEx IGHV dataset for novel alleles with TIgGER detected 271 alleles found in the ComEx dataset as well as four putative additional alleles. IGHV alleles are described here using four naming schemes. Alleles that are present in the IMGT Reference Directory are referred to using IMGT names of the type ‘IGHV1-2*01’. Other non-IMGT alleles that were first reported by Haines and colleagues have been named using a schema producing labels such as ‘J558.1.85’ (35). Non-IMGT sequences that were not named by Haines and colleagues but that appear in the VBASE2 database are referred to by their VBASE2 labels, such as ‘musIGHV021’. Finally, alleles that were first described by Collins and colleagues (19), or were first described in this study have been assigned names such as ‘balbIGHV034’ or ‘b6IGHV040’, according to strain. All confirmed BALB/c IGHV sequences were also ultimately assigned names using the OGRDB Naming Schema (https://github.com/williamdlees/IgLabel, manuscript in preparation).

A novel allele named here as balbIGHV036 was previously reported as a BALB/c IGHV by Corcoran and colleagues (20). A second novel allele appeared to be a variant of balbIGHV027 that differed at the 3’ end of the sequence. No alignments were found to balbIGHV027 itself. This variant allele was also previously reported by Corcoran *et al* (20). It was assigned the name balbIGHV037. A third novel allele had a single difference to balbIGHV009 (balbIGHV009_G32A) and was named balbIGHV038. A truncated version of balbIGHV038 appears as a BALB/c IGHV sequence in the VBASE2 database. The fourth inferred allele was present in the IMGT database but had not been included in ComEx. It was found as IGHV1-12*01_C5T_T6G_T7C_T9A_A12G_T19C_G41A_A139G_G216A_C232T_T239C_A291G_T3 08A and corresponds to IGHV 1S121*01.

When the absence of musIGHV398-utilizing sequences was investigated, a variant with a 7-nucleotide deletion was suspected. It was named balbIGHV039. Investigation of musIGHV269-utilizing sequences highlighted the likely association of this reported IGHV gene with additional 5’ and 3’ nucleotides. A possible allelic variant was also seen. These two sequences were assigned the names b6IGHV040 and balbIGHV041. Re-analysis of the VDJ-C dataset with TIgGER, using an expanded ComEx dataset, identified the presence of balbIGHV039, b6IGHV040 and balbIGHV041.

The six non-IMGT sequences are reported in **Table I**. The seven inferred sequences were added to ComEx, and balbIGHV027 was removed, before a final genotyping and haplotyping analysis was performed.

### Haplotyping identifies a SNP that distinguishes the C57BL/6 and BALB/c *Ighm* CH1 exons

IgM haplotyping began with manual exploration of the IgM constant region exons based on IGHJ1 allele usage. There were 29,593 reads using IGHJ1; 14,071 *01 and 15,522 *03. Alignment of the top IGHJ1*01- and IGHJ1*03-associated *Ighm* sequences to the BALB/c and C57BL/6 genomes confirmed 100% identical matches to the BALBc *Ighm* exons for the *01-associated sequence and 100% identity to the C57BL/6 *Ighm* exons for the *03-associated sequence. A single SNP, rs29176517 (http://www.informatics.jax.org/snp/rs29176517), was identified in the CH1 exon that differentiates IgM from the two strains (see **Table II**). Given the propensity of homopolymer tract errors in PacBio sequencing data from RSII and Sequel I platforms, constant region sequences were indel corrected and the rs29176517 SNP extracted. IGHC calls were assigned to each VDJ-C read using the SNP calls: IgM_B6 (rs29176517a), IgM_BALB (rs29176517g) or ‘uncalled’.

**Table II:**
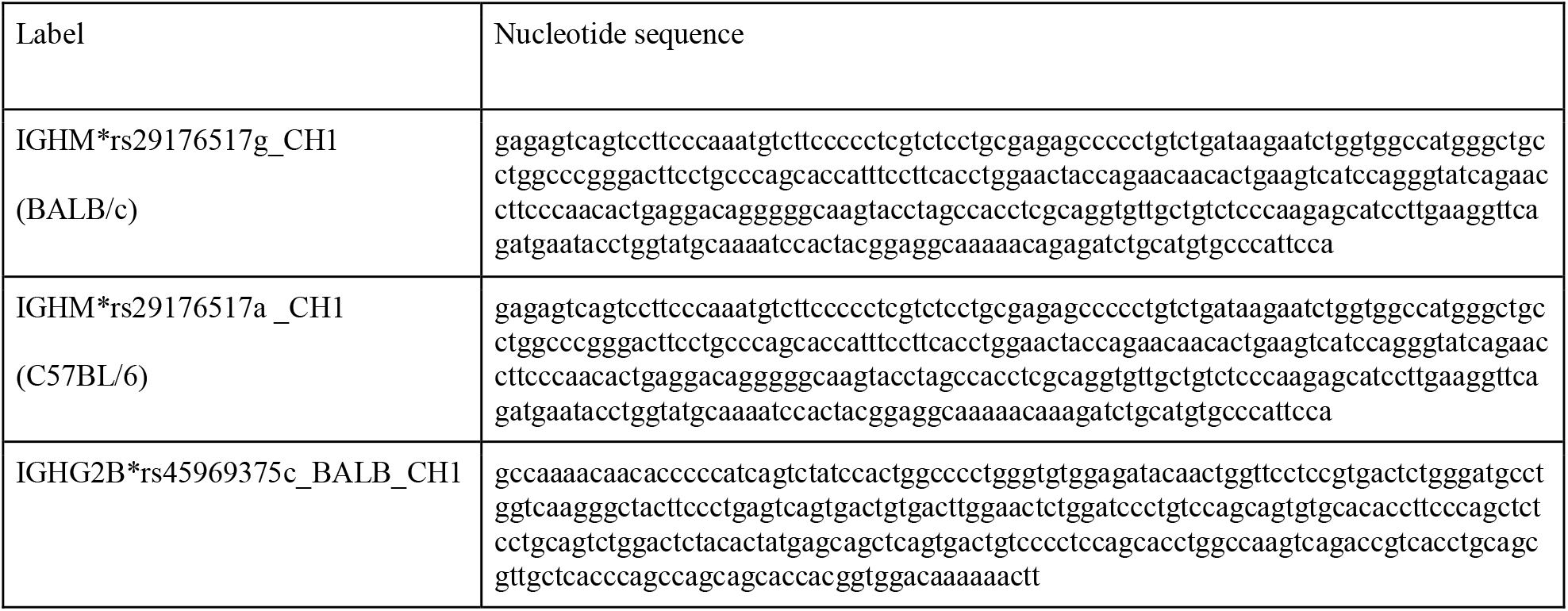
Sequences of inferred BALB/c constant region gene exons used in conjunction with previously reported sequences for C gene-based haplotyping of IGHV genes.

Among the IGHJ1 subset, 85.77% of VDJ-Cs carrying rs29176517g were associated with the BALB/c IGHJ1*01 gene, while 85.98% of VDJ-Cs with rs29176517a were associated with the C57BL/6 IGHJ1*03 gene. This suggests some PCR-based chimerism within the datasets. Overall, there were 58,762 VDJ-Cs matched to the IgM_BALB and 55,445 VDJ-Cs with the IgM_B6.

The C57BL/6 CH1 sequence is identical to an allele that was recently added to the IMGT Reference Directory as IGHM*04. This sequence maps to chr12: 113386037 - 113386351 of the mouse GRCm39/mm39 genome reference sequence. In contrast, the IGHM*01 allele, which is annotated in the IMGT Reference Directory as a C57BL/6 allele, was not observed. The BALB/c CH1 sequence aligns perfectly to chr12: 111920727 – 111921041 of the ‘house mouse 09 May 2016/BALB_cJ_v1’ sequence of the Mouse Genome Project (MGP). This sequence is not in the IMGT Reference Directory and is shown in **Table II**. All other amplified IGHM exons were shared between the two strains, and match sequences in the IMGT Reference Directory.

### Haplotyping identifies SNPs that distinguish C57BL/6 and BALB/c *Ighg2b* genes

The IGHJ1 manual haplotyping approach was applied to the IgG data. The strain specific IgG2a and IgG2c regions were identified along with two IgG2b sequences. Four SNPs were found to distinguish the *Ighg2b* genes between the strains; rs45969375 in the CH1 and rs49934817, rs45822066 and rs46899601 in the CH2. SNP genotyping therefore allowed IgG2b alleles to be assigned to either BALB/c or C57BL/6.

The C57BL/6 IGHG2B*rs45969375t CH1 exon sequence has 100% identity to a new IMGT allele, IGHG2B*03, and to chr12: 113271264 - 113271554 of the mouse GRCm39/mm39 genome reference sequence. The C57BL/6 hinge, CH2 and partial CH3 sequences that were identified here also match IGHG2B*03. The BALB/c IGHG2B* rs45969375c sequence has 100% identity to chr12: 111806470-111806760 of the ‘house mouse 09 May 2016/BALB_cJ_v1’ reference sequence and is shown in **Table I**. It did not match any of the IMGT alleles. The BALB/c hinge, CH2 and partial CH3 sequences are also matches to the IMGT IGHG2B*02 rather than IGHG2B*01 which is annotated as being BALB/c-derived.

BALB/c *Ighg2a* sequences matched perfectly to the IMGT sequence IGHG2A*01 (data not shown). C57BL/6 *Ighg2c* when compared to the IMGT reference directory matched the IGHG2C*01 sequence. The full length of the CH3-CHS exon was not captured by the amplicons and a search of GRCm39 suggests that C57BL/6 CH3-CHS exon matches the IGHG2C*03 sequence (data not shown). The IgG primer was not optimized for *Ighg1* and *Ighg3* associated sequences. Consequently, there was insufficient capture to allow investigation of those genes.

### Genotype and Haplotype inferences using long-read sequencing largely confirm the reported C57BL/6 and BALB/c IGHV genotypes

TIgGER identified 278 IGHV germline alleles in the F1 genotype. 109 known C57BL/6 sequences (IMGT Functionality: 94F, 10P and 5 ORF) were present in the F1 genotype. The pseudogenes were seen as low frequency non-functional rearrangements. We initially confirmed the presence of the musIGHV269 sequence in the C57BL/6 haplotype, and then determined that this sequence is identical to the pseudogene IGHV1-2*01 as it appears in the IMGT Gene Table. This relationship was not identified earlier as IGHV1-2*01 does not appear in the IMGT Reference Set. Closer inspection of the musIGHV269 alignments revealed that both the VBASE2 and IMGT sequences are incomplete. The complete sequence was designated balbIGHV040 (see **Table I**). The full-length sequence is readily identified in the C57BL/6 genome reference sequence. It can also be seen in the IMGT annotations of the genome reference sequence AC073561. Rare reads that represent rearranged sequences of this IGHV allele are also seen in another dataset (ENA: ERR1759753) generated from transcripts derived from C57BL/6 mice (data not shown).

Only two sequences that were previously identified in the C57BL/6 strain were not seen here (IGHV1-74*04 and IGHV8-6*01). These sequences have previously been reported in VDJ repertoires at frequencies of less than 0.1% (19).

Twenty-seven sequences that were identified here, including 14 pseudogenes, were not seen in the previous analysis of the parental strains (19). All but six of them were seen at very low frequencies (<0.01% - 0.03%). This low level of expression could explain their absence from the previous analysis that was undertaken at lower sequencing depth (19). Three of the more abundant sequences were newly inferred alleles that are not present in the IMGT database (see **Table I**).

Nine sequences that IMGT reports as functional C57BL/6 sequences were not seen in either this or previous studies (19): IGHV1-56*01, IGHV1-62-1*01, IGHV1-71*01, IGHV1-74*04, IGHV1S5*01, IGHV5-12-4*01, IGHV6-7*01, IGHV8-6*01 and IGHV8-11*01.

Of the 278 identified IGHV, 169 identified sequences are not present in the C57BL/6-derived GRCm39/mm39 genome reference sequence and are of likely BALB/c origin. This included 17 sequences that were not previously seen (19), and were mostly identified here at very low frequencies (<0.03%). Six sequences that we formerly associated with the strain were not identified. Five of them (balbIGHV016, balbIGHV031, balbIGHV033, IGHV1S75*01 and IGHV1S136*01) were previously seen at frequencies between 0.01% and 0.1% (19). The sixth sequence, musIGHV398, was previously seen at a frequency of 1.56%. Although the musIGHV398 sequence was not seen in the present study, a single full-length, perfect alignment was seen to a sequence that appears to be a pseudogene variant of musIGHV398, with a seven-nucleotide deletion in the FR1 region. This sequence was designated balbIGHV039 (see **Table I**).

The terminal nucleotides of each identified IGHV sequence were subjected to further analysis, as exonuclease processing of the gene ends in VDJ rearrangements can make it difficult to infer the final nucleotides of an IGHV sequence with certainty. Analysis of the frequency distribution of terminal nucleotides at the 3’ ends of each identified BALB/c sequence supported previously reported gene ends in all but a handful of cases. The penultimate nucleotide of the extension of IGHV1S82*01 was shown to be in error, being determined to be G, rather than T. The final nucleotides of IGHV10-1*02 were also shown to be in error, with analysis confirming that the sequence ending is GAGACA, rather than GAGCGA, and is identical to the ending of the IGHV10-1*01 allele. balbIGHV015, musIGHV672, J558-27 and J558-44 may be truncated by 1 or 2 nucleotides, however these sequences are sufficiently distinct from other BALB/c IGHV genes that there was no trouble identifying them in VDJ rearrangements. Finally, the 3’ terminal nucleotides of IGHV13-2*02 (beyond base 320) could not be confirmed, and it is likely that the sequence as described by IMGT is two nucleotides too long.

Haplotype analysis was then conducted using RAbHIT (23), to provide further evidence in support of the strain-specific origins of each gene. To limit the possibility of errors, haplotype analysis was limited to unmutated IgM data. A total of 7,876 VDJ-C sequences were suitable for haplotyping: 3,616 sequences associated with the C57BL/6 IGHM and 4,260 sequences associated with the BALB/c IGHM. Two-hundred and sixty IGHV were successfully assigned to one or both strains, and a representative haplotype plot for genes of the IGHV8 subgroup is shown as **Figure 1**. Eighteen IGHV that were present in the genotype could not be haplotyped. These sequences are listed in **Supplementary Table IV**. In most cases the sequences could not be haplotyped because they were of low frequency and too few reads met the additional haplotype filtering criteria. Seven of these IGHV sequences can be found in the C57BL/6-derived GRCm39/mm39 genome reference sequence, and their identities as rearrangeable C57BL/6 IGHV cannot be doubted. The 11 non-C57BL/6 IGHV are likely BALB/c sequences, but this cannot be confirmed. Seven of the 11 sequences are present in the IMGT reference directory where they are assigned to either the BALB/c strain or the 129 group, and one is the pseudogene variant of musIGHV398 that appears in the BALB/cByJ strain.

**Figure 1:**
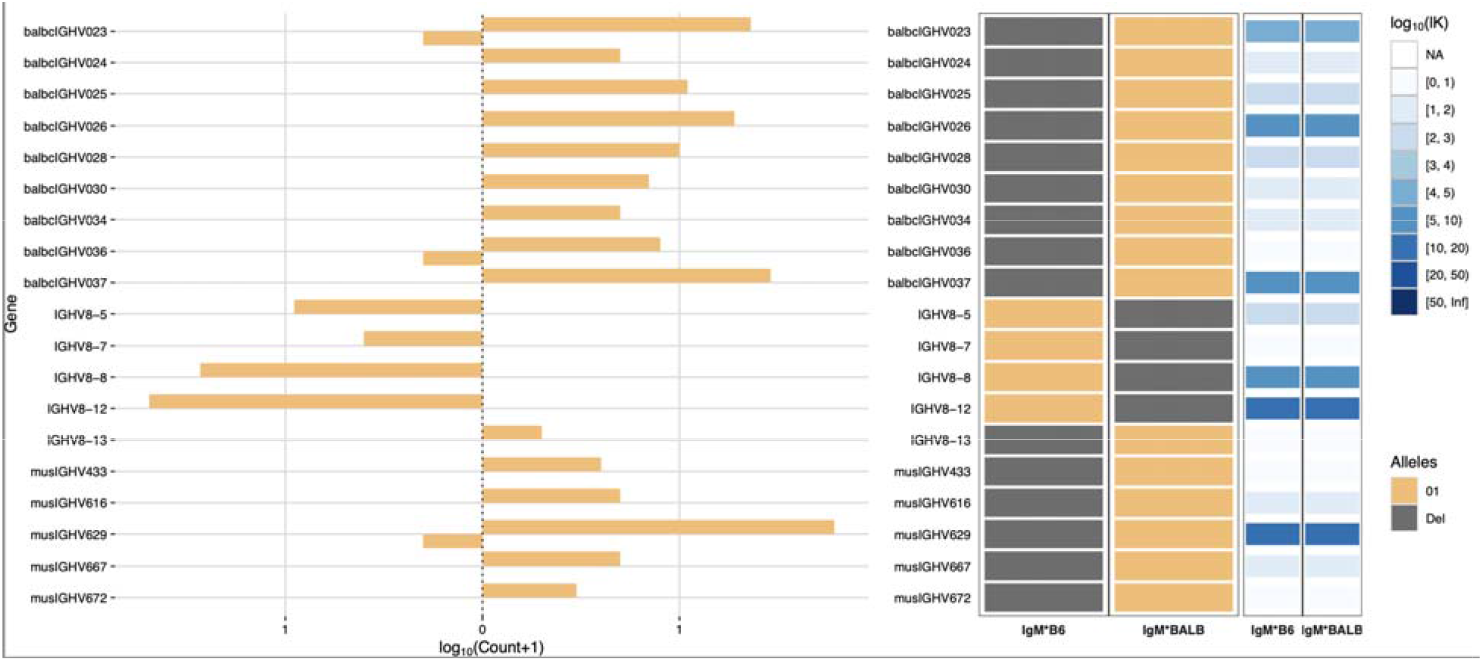
IGHV8 subgroup haplotype plots for BALB/c and C57BL/6. The C57BL/6 chromosome is shown as IgM*B6 and the BALB/c chromosome as IgM*BALB.

Only four sequences (IGHV2-3*01, IGHV2-5*01, IGHV5-2*01 and IGHV5-6*01) were seen in both strains. IGHV1-69*01 was previously reported in both strains (19), but was only confirmed here in C57BL/6 mice. Instead, IGHV1-69*02 was inferred to be present in the BALB/c-derived haplotype.

The strength of the haplotype confirmation process is illustrated by the very low percentage of VDJ-C sequences that apparently included IGHV genes from one strain with IGHM genes from the other, after appropriate filtering to remove sequences for which the IgM allele identification was ambiguous or uncertain. Only 66 (0.84%) of the 7,876 IgM VDJ sequences in the haplotype analysis appear to include chimerism or other problems leading to such haplotyping errors.

### Utilization of IGHV genes varies between F1 and parental strains

The utilization frequencies of the four gene sequences that were confirmed in both strains varied substantially between the parental strains (19), and were different again in the F1 animals. For example, it was previously reported that the IGHV5-6*01 gene was utilized by 0.38% of all C57BL/6 VDJ sequences and the identical IGHV5-6-1*01 gene by 0.27% of BALB/c VDJ sequences. In this study, the sequence was seen in 1.08% of all VDJ rearrangements, including 1.24% of the C57BL/6 chromosomal rearrangements and 0.94% of VDJ of the BALB/c chromosomal rearrangements. Similar increases as well as decreases in utilization were seen for many other sequences. For example, IGHV1S113*01 was previously seen in only 0.01% of BALB/c rearrangements (19), but in F1 animals it was seen in 0.99% of rearrangements of the BALB/c chromosome. In the parental C57BL/6 strain, IGHV1-18*01 was seen in just 0.01% of rearrangements but it was seen in 1.34% of rearrangements of the C57BL/6 chromosome in F1 mice. In contrast, IGHV1-59*01 was seen in 3.40% of all C57BL/6 rearrangements (19), but in the F1 animals it was seen in just 0.41% of rearrangements of the C57BL/6 chromosome. Expression frequencies for each gene in the parental strains and in F1 animals are shown as **Figure 2**.

**Figure 2:**
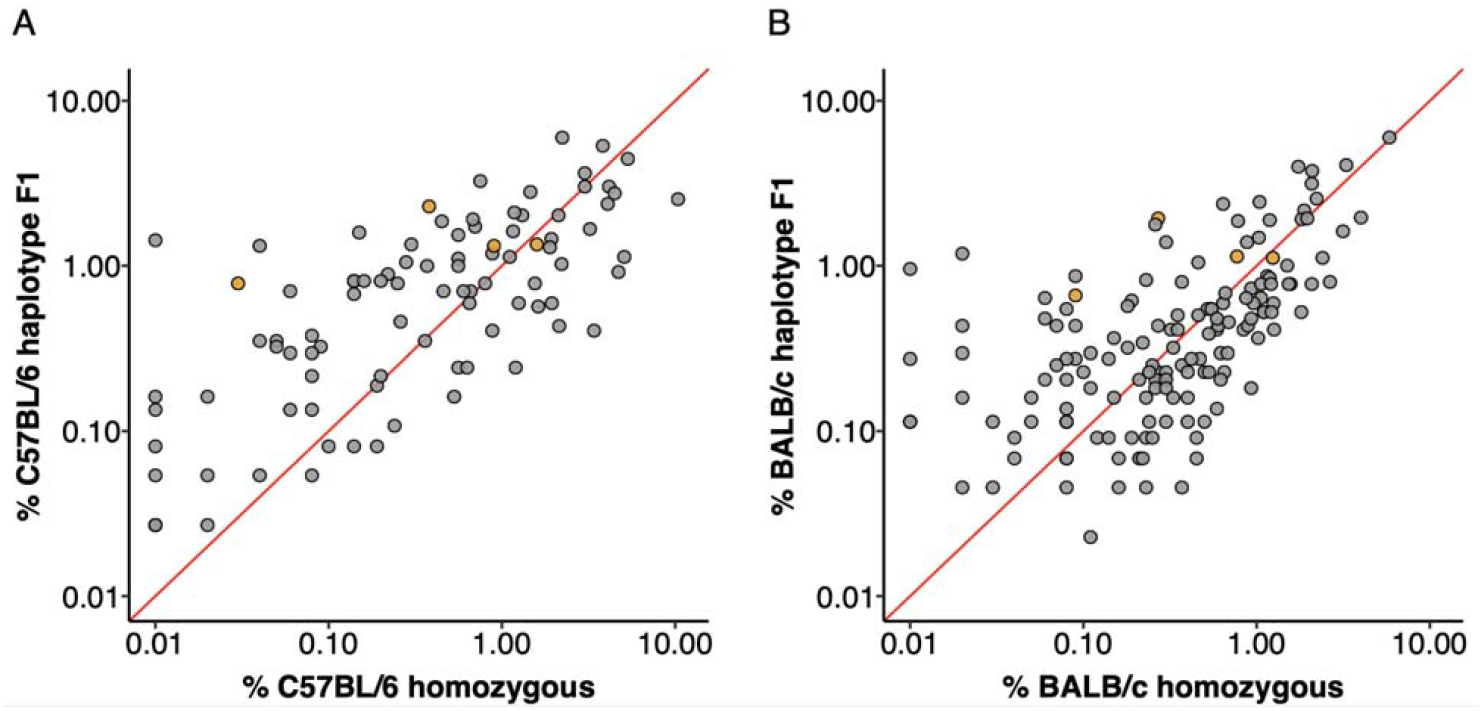
IGHV gene utilization of F1 haplotypes compared to parental strains. Each data point is an IGHV gene, orange filled points are genes shared by the C57BL/6 and BALB/c haplotype. The red line plots y = x and genes with equal utilization in F1 haplotype and homozygous parent strain would fall on this slope. **A)** IGHV gene utilization of C57BL/6 haplotype genes from F1 dataset compared to expression in C57BL/6 homozygous mice. **B)** IGHV gene usage of BALB/c haplotype genes from F1 dataset compared to usage in homozygous BALB/c mice.

### Importance of valid databases for annotation and analysis of hypermutation status

To provide a measure of the errors that can arise from the use of an IGHV Reference Set that poorly represents the IGHV genotype of a mouse strain, we analyzed a dataset of 772,837 BALB/cJ VDJ reads derived from the IgM-encoding transcriptome, documenting the apparent levels of somatic point mutation in each sequence, by counting nucleotide mismatches in the sequence alignments (see **Figure 3**). Although 65.9% of sequences were shown to be unmutated, in analysis using the OGRDB BALB/c Reference Set, only 37.8% of sequences appeared to be unmutated in analysis using the IMGT Reference Set. Differences were conspicuous for sequences with 2, 6, 8, 9, 18 and 19 mismatches. A conspicuous group of sequences was also seen in analysis using the OGRDB Reference Set, with 15 mismatches to the germline alignments. Investigation of these sequences showed them to be almost entirely alignments to the balbIGHV005 sequence. Further investigation showed these sequences to align perfectly to musIGHV398, and the inclusion of this sequence in the Reference Set resulted in 72.1% of sequences being perfect alignments. musIGHV398 was subsequently identified in the MGP BALB/cJ_V1 genome assembly.

**Figure 3:**
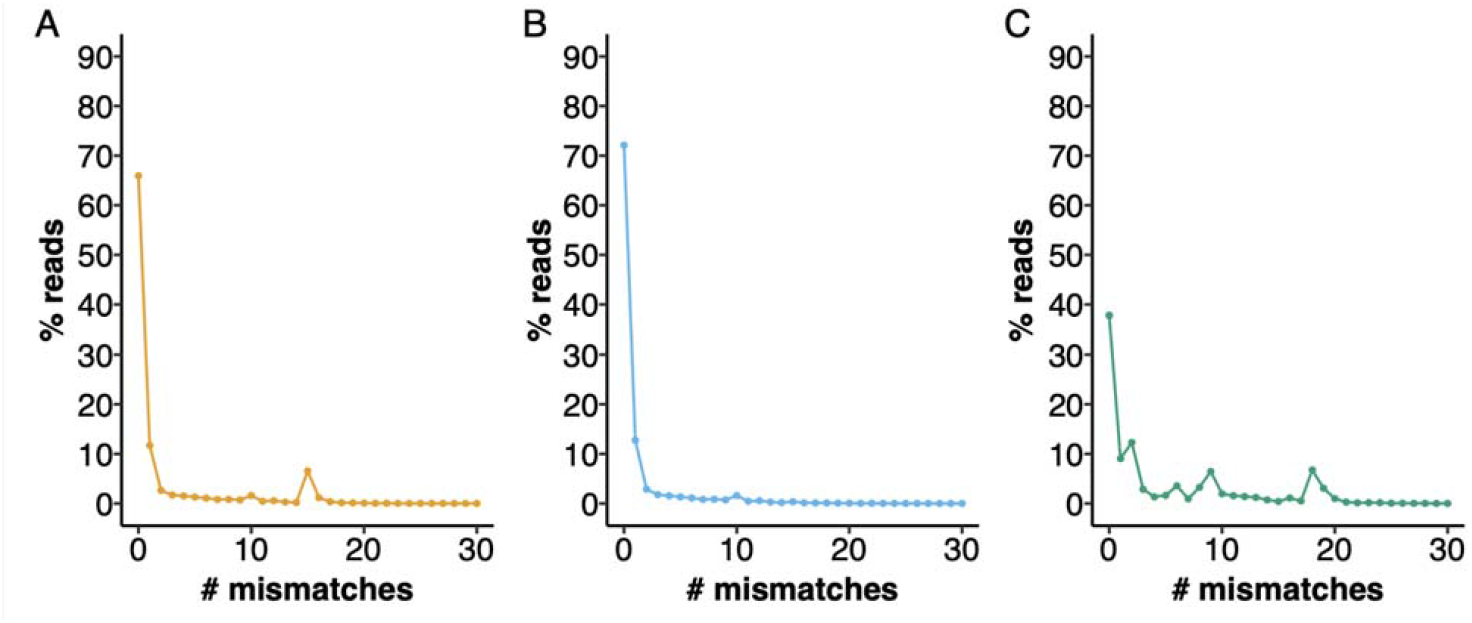
Performance of different IGHV reference sets for the alignment of BALB/c VDJ sequences. IgM reads from BALB/c mice were aligned against different reference directories and the number of mismatches to the closest germline gene was calculated. The percentage of reads with 0 - 30 mismatches are plotted. A) Aligned to OGRDB Reference set without the inclusion of musIGHV398 B) Aligned to OGRDB Reference set with the inclusion of musIGHV398 C) Aligned to IMGT Reference Directory.

The musIGHV398 sequence and its variant (balbIGHV039), as well as the other 162 IGHV sequences that were confirmed as BALB/c genes by haplotype analysis, were used to establish a BALB/c IGHV sequence Reference Set at the OGRDB website (https://ogrdb.airr-community.org/) (24). This set can also be found in the Supplementary Material as the file BALB.xlsx. In addition to their previously-assigned names, the sequences in the BALB/c Reference Set have been given new labels, using the OGRDB Temporary Nomenclature (manuscript in preparation). This Schema recognizes the impossibility of presently determining ‘relationships by descent’ between the genes of the C57BL/6 strain which were used to develop the IMGT nomenclature, and the genes of the BALB/c strain. It is intended that these labels will facilitate the reporting of BALB/c IG sequences until a new community-supported nomenclature can be developed for use with inbred mouse strains.

### Current genomic sequencing does not reliably define IGHV loci content

To investigate the potential to define the IGH loci of inbred mouse strains by annotation of shortread genomic assemblies, we conducted BLAST searches (31) of the Mouse Genome Project (MGP) BALB/cJ_V1 genome assembly (36) for each of the IGHV genes in the OGRDB BALB/c Reference Set. Of the 162 BALB/c sequences, only 70 are present in the MGP assembly. Missing sequences were by no means confined to those sequences that were newly identified in our earlier study. 85 of the sequences are present in the IMGT Reference Directory, but only 37 (43.5%) of these sequences can be found in the MGP assembly. Of the 77 sequences that are not present in the IMGT Reference Directory, only 33 (42.9%) of the sequences are present in the MGP assembly, including 5 of 12 sequences originally reported by Haines and colleagues (35) and 16 of 35 sequences that can be found in the VBASE2 database (37).

## Discussion

Despite the central role of inbred mice in biomedical research, and their historical importance in immunoglobulin gene studies, the germline IGHV genes of inbred mouse strains remain poorly understood. For the IGHV locus, only the genes of the C57BL/6 strain have been fully documented, and this presently limits the use of antibody repertoire analysis in mouse studies.

The Mouse Genome Project has produced genome assemblies of many mouse strains, and analysis of these assemblies might seem like a sensible way to extend the documentation of mouse IGHV genes. Genomic analyses of this kind recently led the IMGT Group to report and name sequences identified in the IGH loci of species including the horse, rat and Rhesus macaque (http://www.imgt.org/IMGTinformation/creations/). There are strong grounds for believing that NGS assemblies of complex genomic regions like the immunoglobulin loci are unlikely to be sufficiently accurate for reliable and comprehensive gene discovery (38, 39). The results of the analysis presented here supports this view. The majority of BALB/c IGHV genes that were identified in this study are not present in the MGP BALB/cJ genome assembly. This is true even for those IGHV sequences that are present within the IMGT Reference Directory, and that are annotated by IMGT as being either BALB/c genes or genes of the highly similar 129S1 strain. The accuracy of NGS assemblies is improving over time, and may well reach the point at which they can be used for receptor gene discovery, however, these results emphasize that their accuracy cannot be assumed without validation against established results and methods.

Germline immunoglobulin gene discovery by inference from AIRR-seq data is an alternative pathway that is well established for the human (23, 40, 41), and confidence in these inferences is often strengthened by haplotype analysis (41, 42). As VDJ rearrangement is a chromosomal event, the association of different IGHV alleles with different human IGHJ6 alleles in IGHJ6-heterozygous individuals allows gene data to be phased. This can lead to the identification of sequences that may appear to be novel alleles, but which arise because of sequence chimerism. Such sequences can easily arise during the PCR amplification of any genes belonging to large sets of highly similar sequences, and it is a problem that has been recognized in studies of antibody genes for decades (27).

It is possible to monitor the extent of the problem within AIRR-seq datasets by careful analysis of gene rearrangements at heterozygous loci. Unfortunately, such loci are not present in inbred mice. An alternative approach identifies chimeric sequences by abnormal clustering of mutations. This approach can also be problematic in the mouse, for it depends upon a thorough knowledge of the germline IGHV gene repertoire of the species. Such analysis is also compromised where multiple genes and their allelic variants are identical or highly similar to one another. This appears to often be the case in inbred mice (22).

We have previously identified IGHV genes in AIRR-seq data generated from inbred BALB/c mice (19, 22). Many of the IGHV genes appeared to rearrange at very low frequencies (0.01% - 0.05%), making their unequivocal identification by inference difficult. It is possible that these inferred IGHV are not real, but rather are chimeric gene products generated by PCR amplifications. In this study, we therefore analyzed VDJ rearrangements from F1 hybrid animals, so that the generation of strain-specific haplotypes could give sufficient confidence in the inferences for the compilation of a BALB/c IGHV Reference Set.

Previous exploration of the C57BL/6 and BALB/c germline repertoire was performed using manual analysis (19), but in recent years, suitable genotyping and haplotyping inference tools have been developed for this purpose (21, 23, 25). Allelic variation is recognized at the mouse IGHJ1 gene. Variation at this locus might be used for IGHV gene haplotype analysis, but the relatively low utilization frequency of the IGHJ1 gene makes it poorly suited to direct IGHJ vs IGHV haplotyping. Genetic variation is also seen within the constant region genes of inbred mice (43). When expressed as VDJ-C genes, however, these constant region (C) genes are too remote from the VDJ sequences to allow haplotype analysis using standard AIRR-Seq sequencing methods (typically based on Illumina’s MiSeq technology). In this study, we therefore used high-throughput single molecule, real time (SMRT) sequencing. Long-read sequencing allowed us to explore allelic variation in genes spanning the entire length of the heavy chain locus, and to perform IGHV gene haplotyping using constant region genes. Against expectations, we identified allelic variation in the CH1 region of the IGHM gene. We also determined that the BALB/c strain carries a previously unreported variant of the *Ighg2b* gene.

The documentation that the BALB/c strain carries an *Ighg2b* constant region gene that differs from that of the C57BL/6 strain will provide researchers with new opportunities for analysis of long-read AIRR-Seq data. In this study, however, we chose to use the heterozygous IGHM locus as the anchor point for haplotype analysis. The general lack of somatic point mutations in the IgM-associated VDJ sequences made the dataset ideal for the documentation of a BALB/c Reference Set by inference.

The results of the haplotype analysis largely confirmed the results of the study of Collins and colleagues (19), though some rarely utilised genes were seen in one study but not the other. This is to be expected, given the sequencing depth of the two studies. Five new BALB/c IGHV genes were identified, and are reported here as balbIGHV036, balbIGHV037, balbIGHV038, balbIGHV039 and balbIGHV041. The agreement between the studies, despite the substantial differences in methodology, demonstrates the reliability of the inference process.

The strain-specific IGHV gene sets defined in this study are comprehensive, but they are still likely to lack some germline genes that are poorly expressed or unexpressed. Some evidence was seen in support of additional sequences - e.g. Genbank IDs: KY199076, KY199111, KY199136 and KY199153 reported from BALB/c studies by Corcoran *et al* (20) - but the evidence was judged to be insufficient to confirm the existence of the genes in this study. Other genes that have apparently been utilized at low frequencies in BALB/c repertoires (19) were not seen here, and have therefore not been included in the OGRDB Reference Sets. Many of these genes may be of questionable functionality, mitigating the impact of their absence from the Reference Set. The IMGT definition of functionality is based upon the presence or absence of key regulatory elements. It therefore may fail to identify the lack of function of genes that encode aberrant nonfunctional polypeptides. Many poorly expressed or unexpressed genes such as IGHV6-7*01, IGHV1-74*04 and IGHV8-6*01 carry unusual codons at critical positions within the sequences. In the human, such unusual codons have been associated with a lack of expression of reportedly functional IGHV genes (44).

Some genes may not have been detected because of differences between the utilization frequencies in the F1 animals, and frequencies previously reported in the parental strains. The expression of antibody genes is relatively predictable. This was first demonstrated in human twin studies (45, 46), and it has also been noted in inbred mice (11). Differences could arise, however, as a result of the genomic context in which the F1 repertoire forms. The naïve antibody repertoire is affected by the processes of negative selection that act to restrict the survival or functionality of self-reactive B cells (47). In F1 animals, the C57BL/6 IGHV gene set and the BALB/c IGHV gene set are each subject to negative selection resulting from both C57BL/6 and BALB/c derived self-antigens. The larger light chain repertoire of F1 animals could also allow those IGHV genes with a tendency to self-reactivity to form more self-tolerant heavy and light chain pairings, thereby increasing the expression of those genes. Some changes in expression frequencies that we observed could also be a consequence of difference in the methodologies of the two studies.

Substrain differences could also explain differences between expression levels reported here for genes such IGHV1S113*01 in the BALB/c mouse and IGHV1-18*01 in the C57BL/6 mouse. The most striking example of a previously-reported IGHV gene that was not seen in this study is the gene musIGHV398. Although it was not called in the F1 genotype, investigation of the absence of musIGHV398-utilizing VDJ in the F1 repertoire led to the identification of a small number of sequences that appeared to utilize a variant of the gene. This variant (balbIGHV039) was subsequently confirmed by genotype analysis. It was also identified in a BAC clone prepared from the BALB/cByJ genome (Kos and Watson: unpublished data). musIGHV398 is present in the MGP BALB/cJ genome assembly and was also seen here in our analysis of BALB/cJ VDJ datasets. It therefore seems that this highly utilized gene defines a critical genetic difference between the BALB/cJ and BALB/cByJ substrains.

Both musIGHV398 and the pseudogene allelic variant balbIGHV039 have been included in the BALB/c Reference Set that is available at the OGRDB website (https://ogrdb.airr-community.org/) (24). Reference Sets for wild-derived strains are also now available at the OGRDB website. These, and the corresponding C57BL/6 Reference Set, will enable better annotation of AIRR-seq studies in comparison to reference germline gene sets that do not take strain differences into account and in some cases even fail to incorporate some highly expressed genes and allelic variants that contribute substantially to the IGHV repertoire.

The BALB/c set that is defined here includes many IGHV genes that are present as truncated sequences in the IMGT reference directory and a large number of genes that have not been named by IMGT or the responsible IUIS Nomenclature Committee. All the BALB/c sequences have been assigned temporary OGRDB labels that will serve adequately until issues relating to the official nomenclature of the mouse can be resolved (48). This resolution will not be possible until careful genomic sequencing allows a proper comparison of the BALB/c and C57BL/6 IGH loci. It will then be possible to determine whether or not there is sufficient correspondence between the two loci for the IMGT C57BL/6-based nomenclature to be applied to the BALB/c strain. The resolution of the assignment of alleles to particular genes is important, for it impacts how some tools handle annotation and analysis. Hopefully, genomic validation of the inference process will also end any uncertainties that might still surround ‘discovery by inference’, and will lead to the exploration of the immunoglobulin genotypes of the many important mouse strains for which high quality genome assemblies are currently unavailable.

## Author Contributions

The study was designed by KJ, CW and AC; library construction and sequencing was performed by CW, JK, WG and ML; data analysis was performed by KJ, MO, WL, CW and AC; manuscript was prepared and edited by KJ, MO, CW, CB, MC, AP, GY and AC.

