## Supplemental Tables I - V for "A BALB/c IGHV Reference Set, defined by haplotype analysis of long-read VDJ-C sequences from F1 (BALB/c / C57BL/6) mice"

Supplementary Table I: Extended sequences of apparently truncated IGHV sequences previously reported as being from the BALB/c or related strains.

| IMGT Gene Name | Full Length Sequence* |
|---|---|
| IGHV1S113*01 | gaggtccagctgcaacagtctggacctgagctggtgaagcctggggcttcagtgaagatatcctgcaagacttctggatacacattcactgaatacaccatgcactgggtgaagcagagccatggaaagagccttgagtggattggaggtattaatcctaacaatggtggtactagctacaaccagaagttcaagggcaaggccacattgactgtagacaagtcctccagcacagcctacatggagctccgcAGCCTGACATCTGAGGATTCTGCAGTCTATTACTGTGCAAGA |
| IGHV1S122*01 | caggtccaactccagcagcctggggctgaactggtgaagcctggggcttcagtgaagttgtcctgcaaggcttctggctacaccttcaccagctactggatgcactgggtgaagctgaggcctggacaaggctttgagtggattggagagattaatcctagcaatggtggtactaactacaatgagaagttcaagagaaaggccacactgactgtagacaaatcctccagcacagcctacatgcaactcagcAGCCTGACATCTGAGGACTCTGCGGTCTATTACTGTACAATA |
| IGHV1S22*01 | CAGGTCCAActgcagcaacctgggtctgagctggtgaggcctggagcttcagtgaagctgtcctgcaaggcttctggctacacattcaccagctactggatgcactgggtgaagcagaggcatggacaaggccttgagtggattggaaatatttatcctggtagtggtagtactaactacgatgagaagttcaagagcaagggcacactgactgtagacacatcctccagcacagcctacatgcacctcagcagcctgacatctgaggactctgcggtctattactgtacaaga |
| IGHV1S33*01 | CAGGTTCAGctgcagcagtctggacctgagctggtgaagcctggggctttagtgaagatatcctgcaaggcttctggttacaccttcacaagctacgatataaactgggtgaagcagaggcctggacagggacttgagtggattggatggatttatcctggagatggtagtactaagtacaatgagaaattcaagggcaaggccacactgactgcagacaaatcctccagcacagcctacatgcagctcagcagcctgacttctgagaactctgcagTCTATTTCTGTGCAAGA |
| IGHV1S34*01 | GAGGTCCAGCTGCAGCAGTCTGGACCTGAGCTAGTGAAGACTGGGGCTTCAGTGAAGatatcctgcaaggcttctggttactcattcactggttactacatgcactgggtcaagcagagccatggaaagagccttgagtggattggatatattagttgttacaatggtgctactagctacaaccagaagttcaagggcaaggccacatttactgtagacacatcctccagcacagcctacatgcagttcaacagcctgacatctgaagactctgcggtctattactgtgcaaga |
| IGHV1S68*02 | caggtccaactgcagcagcctggggctgagcttgtgaagcctgggacttcagtgaagctgtcctgcaaggcttctggctacaacttcaccagctactggataaactgggtgaagctgaggcctggacaaggccttgagtggattggagatatttatcctggtagtggtagtactaactacaatgagaagttcaagagcaaggcc |

| | |
|---|---|
| | acactgactgtagacacatcctccagcacagcctacatgcaactcagcAGCCTGGCATCTGAGGACTCTGCTCTCTATTACTGTGCAAGA |
| IGHV1S72*01 | caggtccaactgcagcagcctggggctgagcttgtgaagcctggggctccagtgaagctgtcctgcaaggcttctggctacaccttcaccagctactggatgaactgggtgaagcagaggcctggacgaggcctcgagtggattggaaggattgatccttccgatagtgaaactcactacaatcaaaagttcaaggacaaggccacactgactgtagacaaatcctccagcacagcctacatccaactcagcAGCCTGACATCTGAGGACTCTGCGGTCTATTACTGTGCAAGA |
| IGHV1S75*01<br><br>** | caggtccaactgcagcagcctggggctgagcttgtgaagcctggggcttcagtgaaaatgtcctgcaaggcttctggctacaccttcaccagctactggataaactgggtgaagcagaggcctggacaaggccttgagtggattggagatatttatcctggtagaggtattactaactacaatgagaagttcaagagcaaggccacactgactctagacacatcctccagcacagcctacatgcagctcagcAGCCTGACATCTGAGGACTCTGCGGTCTATTATTGTTCAAGA |
| IGHV1S82*01<br><br>*** | caggtccaactgcagcagcctggggctgagctggtgaggcctggagcttcagtgaagctgtcctgcaaggcttctggctactccttcaccagctactggatgaactgggtgaagcagaggcctggacaaggccttgagtggattggcatgattcatccttccgatagtgaaactaggttaaatcagaagttcaaggacaaggccacattgactgtagacaaatcctccagcacagcctacatgcaactcagcAGCCCGACATCTGAGGACTCTGCGGTCTATTACTGTGCAAGA |
| IGHV9S8*01<br><br>**** | cagatccagttggtgcagtctggacctgagctgaagaagcctggagagacagtcaagatctcctgcaaggcttctgggtataccttcacaaactatggaatgaactgggtgaagcaggctccaggaaagggtttaaagtggatgggctggataaacactgagactggtgagccaacatatgcagatgacttcaagggacggtttgccttctctttggaaacctctgccagcactgcctatttgcagatcaacaacctcaaaaatgaggacacggctacatatttctgtGCAAGA |
| balbIGHV023 | caggttactctgaaagagtctggccctgggatattgcagccctcccagaccctcagtctgacttgttctttctctgggttttcactgagcacttctggtatgggtgtgagctggattcgtcagccttcaggaaagggtctggagtggctggcacacatttactgggatgatgacaagcgctataacccatccctgaagagccggctcacaatctccaaggatacctccagaaaccaggtattcctcaagatcaccagtgtggacactgcagatactgccacatactactgtgctcgaAGAG |
| balbIGHV026 | caggttactctgaaagagtctggccctgggatattgcagccctcccagaccctcagtctgacttgttctttctctgggttttcactgagcacttctggtatgggtgtaggctggattcgtcagccttcagggaagggtctggagtggctggcacacatttggtgggatgatgacaagcgctataacccagccctgaagagccgactgacaatctccaaggatacctccagcaaccaggtattcctcaagatcgccagtgtggacactgcagatactgccacatactactgtgctcgaataG |

| | |
|---|---|
| balbIGHV028 | caggttactctgaaagagtctggccctgggatattgcagccctcccagaccctcagtctgacttgttctttctctgggttttcactgagcacttctggtatgagtgt<br>aggctggattcgtcagccttcagggaagggtctggagtggctggcacacatttggtggaatgatgataagtactataacccagccctgaaaagccggctcac<br>aatctccaaggatacctccaacaaccaggtattcctcaagatcgccagtgtggtcactgcagatactgccacatactactgtgctcgaataG |
| balbIGHV030 | caggttactctgaaagagtctggccctgggatattgcagccctcccagaccctcagtctgacttgttctttctctgggttttcactgagcacttctggtatgggtgt<br>aggctggattcgtcagccttcaggagagggtctagagtggctggcagacatttggtgggatgacaataagtactataacccatccctgaagagccggctcac<br>aatctccaaggatacctccagcaaccaggtattcctcaagatcaccagtgtggacactgcagatactgccacttactactgtgctcgaagaG |
| balbIGHV032 | caggtccagctgcagcagtctgggcctgagctggtgaggcctggggtctcagtgaagatttcctgcaagggttccggctacacattcactgattatgctatgc<br>actgggtgaagcagagtcatgcaaagagtctagagtggattggagttattagtacttactctggtaatacaaactacaaccagaagtttaagggcaaggccac<br>aatgactgtagacaaatcctccagcacagcctatatggaacttgccagattgacatctgaggattctgccatctattactgtGCAAGA |
| balbIGHV034 | Caggttactctgaaagagtctggccctgggatattgcagccctcccagaccctcagtctgacttgttctttctctgggttttcactgagcacttctggtatgggtgt<br>aggctggattcgtcagccttcagggaagggtctggagtggctggcacacatttggtgggatgatgataagtactataacacagccctgaagagcgggctca<br>caatctccaaggatacctccaaaaaccaggtcttcctcaagatcgccagtgtggacactgcagatactgccacatactactgtgctcgaataG |
| J558.27 | caggtccagctgcagcagtctggacctgagctggtgaagcctggggcttcagtgaagatgtcctgcaaggcttctggctacaccttcacaagctactatatac<br>actgggtgaagcagaggcctggacagggacttgagtggattggatggatttatcctggagatggtagtactaagtacaatgagaagttcaagggcaagacc<br>acactgactgcagacaaatcctccagcacagcctacatgttgctcagcagcctgacctctgaggactctgcgatctatttctgtgcaagA |
| J558.44 | caggtccagctgcagcagtctggggctgaactggcaagacctggggcctcagtgaagatgtcctgcaaggcttctggctacacctttactagctacacgatg<br>cactgggtaaaacagaggcctggacagggtctggaatggattggatacattaatcctagcagtggttatactaattacaatcagaagttcaaggacaaggcca<br>cattgactgcagacaaatcctccagcacagcctacatgcaactgagcagcctgacatctgaggactctgcagtctattactgtgcaaGA |
| musIGHV021 | caggttcagctccagcagtctggggctgagctggcaagacctggggcttcagtgaagttgtcctgcaaggcttctggctacacctttactagctactggatgc<br>agtgggtaaaacagaggcctggacagggtctggaatggattggggctatttatcctggagatggtgatactaggtacactcagaagttcaagggcaaggcc<br>acattgactgcagataaatcctccagcacagcctacatgcaactcagcagcttggcatctgaggactctgcggtctattactgtGCAAGA |
| musIGHV616 | caggttactctgaaagagtctggccctgggatattgcagccctcccagaccctcagtctgacttgttctttctctgggttttcactgagcacttctaatatgggtgt<br>aggctggattcgtcagccttcagggaagggtctggagtggctgttacacattttgtggaatgatagtaagtactataacccagccctgaagagccggctcaca<br>atctccaaggatacctacaacaaccaggtattcctcaagatcgccaatgtggacactgcagatactgccacatactactgtgctcgaATAG |

| | |
|---|---|
| musIGHV672 | caggttattctgaaagagtctggccctggaatattgcagccctctcagaccctcagtctgacttgttctttctctgggttttcacttagcacttatggtacagctgtg aactggattcgtcagccttcaggaaagggtctggagtggttggcacaaattgggtcagatgatagcaagctctataacccatttctgaaaagccgaatcacaat ctccaaggatacctccaacagccaggtattcctcaagatcactagtgtggacactgaagattctgccacatactactgtgctaacAGA |

* Extensions are shown in Uppercase
** Not confirmed in this study
*** The extension originally ended in the terminal nucleotides TA, but were determined to be GA by analysis of the rearranged gene ends.
**** Unable to be confirmed by haplotype analysis

Supplementary Table II: Mouse IGHV sequences in the IMGT Reference Directory that have been changed since the analysis by Collins *et al* (Collins et al., 2015), with altered or added sequences in uppercase.

| | |
|---|---|
| IGHV2-9*02<br><br>(previously named IGHV2-9-1*01) | caggtgcagctgaaggagtcaggacctggcctggtggcgccctcacagagcctgtccatcacttgcactgtctctgggttttcattaaccagctat<br>ggtgtacactgggttcgccagcctccaggaaagggtctggagtggctgggagtaatatgggctggtggaagcacaaattataattcggctctcat<br>gtccagactgagcatcagcaaagacaactccaagagccaagttttcttaaaaatgaacagtctgcaaactgatgacacagccatgtactactgtgc<br>cagaga |
| IGHV2-9-1*01<br><br>(previously musIGHV211) | caggtgcagctgaaggagtcaggacctggcctggtggcgccctcacagagcctgtccatcacatgcactgtctctgggttctcattaaccagctat<br>gctataagctgggttcgccagccaccaggaaagggtctggagtggcttggagtaatatggactggtggaggcacaaattataattcagctctcaa<br>atccagactgagcatcagcaaagacaactccaagagtcaagttttcttaaaaatgaacagtctgcaaactgatgacacagccaggtactactgtgc<br>cagaAA |
| IGHV2-9-2*01<br><br>(previously IGHV2-9*02) | caggtgcaactgaaggagtcaggacctggcctggtggcgccctcacagagcctgtccattacctgcactgtctctgggttctcattaaccagctat<br>gatataagctggattcgccagccaccaggaaagggtctggagtggcttggagtaatatggactggtggaggcacaaattataattcagctttcatg<br>tccagactgagcatcagcaaggacaactccaagagccaagttttcttaaaaatgaacagtctgcaaactgatgacacagccatatattactgtgtaa<br>gaga |
| IGHV4-1*01 | gaggtgaagcttctccagtctggaggtggcctggtgcagcctggaggatccctgaaactctcctgtgcagcctcaggaatcgattttagtagatac<br>tggatgagttgggttcggcgggctccagggaaaggactagaatggattggagaaattaatccagatagcagtacaataaactatgcaccatctcta<br>aaggataaattcatcatctccagagacaacgccaaaaatacgctgtacctgcaaatgagcaaagtgagatctgaggacacagccctttattactgt<br>gcaagaCC |
| IGHV5-6-5*01<br><br>(previously IGHV5S12*01) | gaagtgaagctggtggagtctgggggaggcttagtgaagcctggagggtccctgaaactctcctgtgcagcctctggattcactttcagtagctat<br>gccatgtcttgggttcgccagactccagagaagaggctggagtgggtcgcatccattagtagtggtggtagcacctactatccagacagtgtgaa<br>gggccgattcaccatctccagagataatgccaggaacatcctgtacctgcaaatgagcagtctgaggtctgaggacacggccatgtattactgtg<br>caagagg |
| IGHV5-15*01 | gaggtgaagctggtggagtctgggggaggcttagtgcagcctggagggtccctgaaactctcctgtgcagcctctggattcactttcagtgactac<br>ggaatggcgtgggttcgacaggctccaaggaaggggcctgagtgggtagcattcattagtaatttggcatatagtatctactatgcagacactgtg |

| | |
|---|---|
| | acgggccgattcaccatctctagagagaatgccaagaacaccctgtacctggaaatgagcagtctgaggtctgaggacacggccatgtattactg tgcaagaCA |
| IGHV5-16*01 | gaagtgaagctggtggagtctgagggaggcttagtgcagcctggaagttccatgaaactctcctgcacagcctctggattcactttcagtgactatt acatggcttgggtccgccaggttccagaaaagggtctagaatgggttgcaaacattaattatgatggtagtagcacctactatctggactccttgaa gagccgtttcatcatctcgagagacaatgcaaagaacattctatacctgcaaatgagcagtctgaagtctgaggacacagccacgtattactgtgc aagaGA |
| IGHV7-1*01 | gaggtgaagctggtggaatctggaggaggcttggtacagtctgggcgttctctgagactctcctgtgcaacttctgggttcaccttcagtgatttcta catggagtgggtccgccaagctccagggaagggactggagtggattgctgcaagtagaaacaaagctaatgattatacaacagagtacagtgc atctgtgaagggtcggttcatcgtctccagagacacttcccaaagcatcctctaccttcagatgaatgccctgagagctgaggacactgccatttatt actgtgcaagaGATGCA |
| IGHV8-6*01 | caggttactctgaaagagtctggccctggtatattgcagccctcgcagaccctcagtctgacttgttctttctctgggttttcactgagtacttttggtat gggtgtgagctggattcgtcagccttcagggaaggatctggagtggctggcacacatttattgggatgatgacaagcactataacccatccttgaa gagccagctcagaatctccaaggatacctccaacaaccaggtattcctcaagatcaccactgtggacactgtagatactgccacatactactgtgc tcgaAGAG |
| IGHV10-3*01 | gaggtgcagcttgttgagtctggtggaggattggtgcagcctaaaggatcattgaaactctcatgtgccgcctctggtttcaccttcaatacctatgc catgcactgggtccgccaggctccaggaaagggtttggaatgggttgctcgcataagaagtaaaagtagtaattatgcaacatattatgccgattc agtgaaagacagattcaccatctccagagatgattcacaaagcatgctctatctgcaaatgaacaacctgaaaactgaggacacagccatgtatta ctgtgtgagagA |

Supplementary Table III: Constant Region Genes inferred from VDJ-C datasets.

| Label | Exon | Strain | Source |
|---|---|---|---|
| IGHG2A*01 | CH1 | BALB/c | IMGT |
| IGHG2A*01 | Hinge | BALB/c | IMGT |
| IGHG2A*01 | CH2 | BALB/c | IMGT |
| IGHG2A*01 | CH3-CHS | BALB/c | IMGT |
| IGHG2B*rs45969375c | CH1 | BALB/c | Inferred |
| IGHG2B*03 (rs45969375t) | CH1 | C57BL/6 | IMGT[a] |
| IGHG2B*02/03 (identical) | Hinge | Both | IMGT |
| IGHG2B*02 | CH2 | BALB/c | IMGT |
| IGHG2B*03 | CH2 | C57BL/6 | IMGT[a] |
| IGHG2C | CH1 | C57BL/6 | IMGT |
| IGHG2C*01 | Hinge | C57BL/6 | IMGT |
| IGHG2C*01 | CH2 | C57BL/6 | IMGT |
| IGHG2C*03 | CH3-CHS | C57BL/6 | IMGT[b] |
| IGHM*rs29176517g | CH1 | BALB/c | Inferred |
| IGHM*04 (rs29176517a) | CH1 | C57BL/6 | IMGT[a] |

| | | | |
|---|---|---|---|
| IGHM*02 | CH2 | Both | IMGT |
| IGHM*02 | CH3 | Both | IMGT |

[a]The sequence can be found in the IMGT database, but is not present in the IMGT Mouse Gene Table. Its association with the strain was confirmed in this study using IGHJ1-based haplotyping of VDJ-C data.

[b]The sequence is available in the IMGT Mouse Gene Table, but is incorrectly assigned. Its association with the strain was confirmed by alignment with the Mouse Genome Reference Sequence.

Supplementary Table IV: IGHV genes identified in the F1 mouse genotype that were unable to be haplotyped.

| | Present in the mouse GRCm39/mm39 genome reference sequence | Present in the BALB_cJ_v1' sequence of the Mouse Genome Project (MGP) |
|---|---|---|
| IGHV1-14*01 | Y | N |
| IGHV1-70*01 | Y | N |
| IGHV1-79*01 | Y | N |
| IGHV2-6-6*01 | N | Y |
| IGHV3-3*02 | N | Y |
| IGHV3-4*02 | N | Y |
| IGHV5-1*02 | N | Y |
| IGHV6-4*01 | Y | N |
| IGHV6-4*02 | N | N |
| IGHV1-34*01 | Y | N |
| IGHV1-37*01 | Y | N |
| IGHV8-4*01 | Y | N |
| IGHV004b*01 | N | N |

| | | |
|---|---|---|
| IGHV014b*01 | N | N |
| IGHV017b*01 | N | Y |
| IGHV1S132*01 | N | Y |
| IGHV9S8*01 | N | N |
| balbIGHV039 | N | N |

Supplementary Table V: The OGRDB BALB/c Reference Set Version 1 (25/2/2022), showing OGRDB labels, IMGT names and previously-assigned non-IMGT names.

| **Label** | **IMGT Name** | **Alt Names** | | **Sequence** |
|---|---|---|---|---|
| IGHV-22XF | | Haines: J558.44 | F | CAGGTCCAGCTGCAGCAGTCTGGGGCTGAACTGGCAAGACCTGGGGCCTCAGTGAAGATGTCCTGCA<br>AGGCTTCTGGCTACACCTTTACTAGCTACACGATGCACTGGGTAAAACAGAGGCCTGGACAGGGTCTG<br>GAATGGATTGGATACATTAATCCTAGCAGTGGTTATACTAATTACAATCAGAAGTTCAAGGACAAGGC<br>CACATTGACTGCAGACAAATCCTCCAGCACAGCCTACATGCAACTGAGCAGCCTGACATCTGAGGACT<br>CTGCAGTCTATTACTGTGCAAGA |
| IGHV-2KUO | | VBASE2: musIGHV612 | F | CAGGGTCAGATGCAGCAGTCTGGAGCTGAGCTGGTGAAGCCTGGGGCTTCAGTGAAGCTGTCCTGCA<br>AGACTTCTGGCTTCACCTTCAGCAGTAGCTATATAAGTTGGTTGAAGCAAAAGCCTGGACAGAGTCTT<br>GAGTGGATTGCATGGATTTATGCTGGAACTGGTGGTACTAGCTATAATCAGAAGTTCACAGGCAAGG<br>CCCAACTGACTGTAGACACATCCTCCAGCACAGCCTACATGCAATTCAGCAGCCTGACAACTGAGGAC<br>TCTGCCATCTATTACTGTGCAAGA |
| IGHV-2UIT | | JACKSON: balbIGHV015 | F | CAGGTCCAACTGCAGCAGCCTGGGGCTGAGCTGGTGAGGCCTGGGGCTTCAGTGAAGCTGTCCTGCA<br>AGGCTTCTGGCTACACGTTCACCAGCTACTGGATGAACTGGGTTAAGCAGAGGCCTGAGCAAGGCCTT<br>GAGTGGATTGGAAGGATTGATCCTTACGATAGTGAAACTCACTACAATCAAAAGTTCAAGGACAAGG<br>CCATATTGACTGTAGACAAATCCTCCAGCACAGCCTACATGCAACTCAGCAGCCTGACATCTGAGGACT<br>CTGCGGTCTATTACTGTGCAAGA |
| IGHV-2UQD | | VBASE2: musIGHV398 | F | GAGGTCCAGCTGCAACAGTCTGGACCTGAGCTGGTGAAGCCTGGAGCTTCAATGAAGATATCCTGCA<br>AGGCTTCTGGTTACTCATTCACTGGCTACACCATGAACTGGGTGAAGCAGAGCCATGGAAAGAACCTT<br>GAGTGGATTGGACTTATTAATCCTTACAATGGTGGTACTAGCTACAACCAGAAGTTCAAGGGCAAGGC<br>CACATTAACTGTAGACAAGTCATCCAGCACAGCCTACATGGAGCTCCTCAGTCTGACATCTGAGGACTC<br>TGCAGTCTATTACTGT |
| IGHV-32WJ | | VBASE2: musIGHV629 | F | CAGGTTACTCTGAAAGAGTCTGGCCCTGGGATATTGCAGCCCTCCCAGACCCTCAGTCTGACTTGTTCT<br>TTCTCTGGGTTTTCACTGAGCACTTCTGGTATGGGTGTGAGCTGGATTCGTCAGCCTTCAGGAAAGGG<br>TCTGGAGTGGCTGGCACACATTTACTGGGATGATGACAAGCGCTATAACCCATCCCTGAAGAGCCGGC<br>TCACAATCTCCAAGGATACCTCCAGCAACCAGGTATTCCTCAAGATCACCAGTGTGGACACTGCAGATA<br>CTGCCACATACTACTGTGCTCGAAGAG |
| IGHV-37IE | IGHV5-17*02 | | F | GATGTGCAGCTGGTGGAGTCTGGGGGAGGCTTAGTGCAGCCTGGAGGGTCCCGGAAACTCTCCTGTG<br>CAGCCTCTGGATTCACTTTCAGTAGCTTTGGAATGCACTGGGTTCGTCAGGCTCCAGAGAAGGGGCTG<br>GAGTGGGTCGCATACATTAGTAGTGGCAGTAGTACCATCTACTATGCAGACACAGTGAAGGGCCGAT |

| | | | | |
|---|---|---|---|---|
| | | | | TCACCATCTCCAGAGACAATCCCAAGAACACCCTGTTCCTGCAAATGACCAGTCTAAGGTCTGAGGAC<br>ACGGCCATGTATTACTGTGCAAGA |
| IGHV-3AN3 | IGHV9-2*02 | | F | CAGATCCAGTTGGTGCAGTCTGGACCTGAGCTGAAGAAGCCTGGAGAGACAGTCAAGATCTCCTGCA<br>AGGCTTCTGGCTATACCTTCACAAACTATGCAATGCACTGGGTGAAGCAGGCTCCAGGAAAGGGTTTA<br>AAGTGGATGGGCTGGAAATACACCAACACTGGAGAGCCAACATATGGTGATGACTTCAAGGGACGGT<br>TTGCCTTCTCTTTGGAAACCTCTGCCAGCACTGCCTATTTGCAGATCAACAACCTCAAAAATGAGGACA<br>TGGCTACATATTTCTGTGCAAGA |
| IGHV-3GCO | | VBASE2: musIGHV014 | F | CAGGTCCAGTTGCAGCAGTCTGGAGCTGAGCTGGTAAGGCCTGGGACTTCAGTGAAGATATCCTGCA<br>AGGCTTCTGGCTACACCTTCACTAACTACTGGCTAGGTTGGGTAAAGCAGAGGCCTGGACATGGACTT<br>GAGTGGATTGGAGATATTTACCCTGGAGGTGGTTATACTAACTACAATGAGAAGTTCAAGGGCAAGG<br>CCACACTGACTGCAGACACATCCTCCAGCACTGCCTACATGCAGCTCAGTAGCCTGACATCTGAGGACT<br>CTGCTGTCTATTTCTGTGCAAGA |
| IGHV-3GZS | IGHV1S135*01 | | F | GAGATCCAGCTGCAGCAGTCTGGACCTGAGCTGGTGAAGCCTGGGGCTTCAGTGAAGGTATCCTGCA<br>AGGCTTCTGGTTACTCATTCACTGACTACAACATGTACTGGGTGAAGCAGAGCCATGGAAAGAGCCTT<br>GAGTGGATTGGATATATTGATCCTTACAATGGTGGTACTAGCTACAACCAGAAGTTCAAGGGCAAGGC<br>CACATTGACTGTTGACAAGTCCTCCAGCACAGCCTTCATGCATCTCAACAGCCTGACATCTGAGGACTC<br>TGCAGTCTATTACTGTGCAAGA |
| IGHV-3QFD | IGHV14-4*02 | | F | GAGGTTCAGCTGCAGCAGTCTGGGGCAGAGCTTGTGAGGTCAGGGGCCTCAGTCAAGTTGTCCTGCA<br>CAGCTTCTGGCTTCAACATTAAAGACTACTATATGCACTGGGTGAAGCAGAGGCCTGAACAGGGCCTG<br>GAGTGGATTGGATGGATTGATCCTGAGAATGGTGATACTGAATATGCCCCGAAGTTCCAGGGCAAGG<br>CCACTATGACTGCAGACACATCCTCCAACACAGCCTACCTGCAGCTCAGCAGCCTGACATCTGAGGAC<br>ACTGCCGTCTATTACTGTAATGCA |
| IGHV-3QLH | | Haines: J558.18 | F | CAGGTCCAGCTGCAGCAGTCTGGACCTGAGCTGGTGAAGCCTGGGGCCTCAGTGAAGATTTCCTGCA<br>AAGCTTCTGGCTACGCATTCAGTAGCTCTTGGATGAACTGGGTGAAGCAGAGGCCTGGACAGGGTCTT<br>GAGTGGATTGGACGGATTTATCCTGGAGATGGAGATACTAACTACAATGGGAAGTTCAAGGGCAAGG<br>CCACACTGACTGCAGACAAATCCTCCAGCACAGCCTACATGCAGCTCAGCAGCCTGACCTCTGTGGAC<br>TCTGCGGTCTATTTCTGTGCAAGA |
| IGHV-47EU | IGHV1S12*01 | | F | CAGGTCCAGCTGCAGCAATCTGGACCTGAGCTGGTGAAGCCTGGGGCTTCAGTGAAGATATCCTGCA<br>AGGCTTCTGGCTATACCTTCACAAGCTACTATATACACTGGGTGAAGCAGAGGCCTGGACAGGGCCTT<br>GAGTGGATTGGATATATTTATCCTAGAGATGGTAGTACTAATTACAATGAGAAGTTCAAGGGCAAGGC<br>CACACTGACTGCAGACACATCCTCCAGCACAGCCTACATGCAGCTCAGCAGCCTGACATCTGAGGACT<br>CTGCAGTCTATTTCTGTGCAAGA |

| | | | | |
|---|---|---|---|---|
| IGHV-4FMT | IGHV5-15*02 | | F | GAGGTGAAGCTGGTGGAGTCTGGGGGAGGCTTAGTGCAGCCTGGAGGGTCCCGGAAACTCTCCTGT<br>GCAGCCTCTGGATTCACTTTCAGTGACTACGGAATGGCGTGGGTTCGACAGGCTCCAG GGAAGGGGC<br>CTGAGTGGGTAGCATTCATTAGTAATTTGGCATATAGTATCTACTATGCAGACACTGTGACGGGCCGA<br>TTCACCATCTCTAGAGAGAATGCCAAGAACACCCTGTACCTGGAAATGAGCAGTCTGAGGTCTGAGGA<br>CACAGCCATGTACTACTGTGCAAGGGA |
| IGHV-4L6T | | VBASE2: musIGHV577 | F | CAGGTCCAACTGCAGCAGCCTGGGGCTGAACTGGTGAAGCCTGGGACTTCAGTGAAAATGTCCTGCA<br>AGGCTTCTGGCTACACCTTCACCAGCTACTGGATGCACTGGGTGAAGCAGAGGCCGGGACAAGGCCT<br>TGAGTGGATTGGAGATATTTATCCTGGTAGTGATAGTACTAACTACAATGAGAAGTTCAAGAGCAAGG<br>CCACACTGACTGTAGACACATCCTCCAGCACAGCCTACATGCAACTCAGCAGCCTGACATCTGAGGACT<br>CTGCGGTCTATTACTGTGCAAGA |
| IGHV-4T4X | | VBASE2: musIGHV042 | F | CAGGTCCAGCTGCAGCAGTCTGGGGCTGAGCTGGTGAGGCCTGGGGCCTCAGTGAAGATTTCCTGCA<br>AGGCTTTTGGCTACACCTTCACAAACCATCATATAAACTGGGTGAAGCAGAGGCCTGGACAGGGCCTG<br>GACTGGATTGGATATATTAATCCTTATAATGATTATACTAGCTACAACCAGAAGTTCAAGGGCAAGGC<br>CACATTGACTGTAGACAAATCCTCCAGCACAGCCTATATGGAGCTTAGCAGCCTGACATCTGAGGACT<br>CTGCAGTCTATTACTGTGCAAGA |
| IGHV-4X2I | IGHV11-2*02 | | F | GAAGTGCAGCTGTTGGAGACTGGAGGAGGCTTGGTGCAACCTGGGGGGTCACGGGGACTCTCTTGT<br>GAAGGCTCAGGGTTTACTTTTAGTGGCTTCTGGATGAGCTGGGTTCGACAGACACCTGGGAAGACCCT<br>GGAGTGGATTGGAGACATTAATTCTGATGGCAGTGCAATAAACTACGCACCATCCATAAAGGATCGAT<br>TCACTATCTTCAGAGACAATGACAAGAGCACCCTGTACCTGCAGATGAGCAATGTGCGATCTGAGGAC<br>ACAGCCACGTATTTCTGTATGAGATA |
| IGHV-4YZL | | VBASE2: musIGHV394 | F | GAGGTCCAGCTGCAACAATCTGGACCTGAGCTGGTGAAGCCTGGGGCTTCAGTGAAGATGTCCTGTA<br>AGGCTTCTGGATACACATTCACTGACTACTACATGAAGTGGGTGAAGCAGAGTCATGGAAAGAGCCTT<br>GAGTGGATTGGAGATATTAATCCTAACAATGGTGGTACTAGCTACAACCAGAAGTTCAAGGGCAAGG<br>CCACATTGACTGTAGACAAATCCTCCAGCACAGCCTACATGCAGCTCAACAGCCTGACATCTGAGGACT<br>CTGCAGTCTATTACTGTGCAAGA |
| IGHV-56YB | | JACKSON: balbIGHV025 | F | CAGGTTACTCTGAAAGAGTCTGGCCCTGGGATATTGCAGCCCTCCCAGACCCTCAGTCTGACTTGTTCT<br>TTCTCTGGGTTTTCACTGAGCACTTCTGGTATGGGTGTAGGCTGGATTCGTCAGCCATCAGGGAAGGG<br>TCTGGAGTGGCTGGCACACATTTGGTGGGATGATGTCAAGCGCTATAACCCAGCCCTGAAGAGCCGA<br>CTGACTATCTCCAAGGATACCTCCAGCAGCCAGGTATTCCTCAAGATCGCCAGTGTGGACACTGCAGA<br>TACTGCCACATACTACTGTGCTCGAATAG |
| IGHV-5CCQ | IGHV11-1*02 | | F | GAAGTGCAGCTGTTGGAGACTGGAGGAGGCTTGGTGCAACCTGGGGGGTCACGGGGACTCTCTTGT<br>GAAGGCTCAGGGTTCACTTTTAGTGGCTTCTGGATGAGCTGGGTTCGACAGACACCTGGGAAGACCG<br>TGGAGTGGATTGGAGACATTAATTCTGATGGCAGTGCAATAAACTACGCACCATCCATAAAGGATCGA |

| | | | | |
|---|---|---|---|---|
| | | | | TTCACTATCTTCAGAGACAATGACAAGAGCACCCTGTACCTGCAGATGAGCAATGTGCGATCTGAGGA<br>CCCAGCCACGTATTTCTGTATGAGATA |
| IGHV-5F5R | IGHV5-6-5*01 | | F | GAAGTGAAGCTGGTGGAGTCTGGGGGAGGCTTAGTGAAGCCTGGAGGGTCCCTGAAACTCTCCTGTG<br>CAGCCTCTGGATTCACTTTCAGTAGCTATGCCATGTCTTGGGTTCGCCAGACTCCAGAGAAGAGGCTG<br>GAGTGGGTCGCATCCATTAGTAGTGGTGGTAGCACCTACTATCCAGACAGTGTGAAGGGCCGATTCAC<br>CATCTCCAGAGATAATGCCAGGAACATCCTGTACCTGCAAATGAGCAGTCTGAGGTCTGAGGACACG<br>GCCATGTATTACTGTGCAAGA |
| IGHV-5ISV | | VBASE2: musIGHV616 | F | CAGGTTACTCTGAAAGAGTCTGGCCCTGGGATATTGCAGCCCTCCCAGACCCTCAGTCTGACTTGTTCT<br>TTCTCTGGGTTTTCACTGAGCACTTCTAATATGGGTGTAGGCTGGATTCGTCAGCCTTCAGGGAAGGG<br>TCTGGAGTGGCTGTTACACATTTTGTGGAATGATAGTAAGTACTATAACCCAGCCCTGAAGAGCCGGC<br>TCACAATCTCCAAGGATACCTACAACAACCAGGTATTCCTCAAGATCGCCAATGTGGACACTGCAGATA<br>CTGCCACATACTACTGTGCTCGAATAG |
| IGHV-5LN7 | | VBASE2: musIGHV692 | F | GAGGTCCAGCTGCAGCAGTCTGGACCTGAGCTGGTGAATCCAGTGGCTTCAGTGAAGATATCCTGCA<br>AGGCTTCTGGTTACTCATTCACTGGCTACTACATACACTGGGTGAAGCAGGGTCCTAGAAAGAGCCTT<br>GAGTGGATTGGATATATTAGTTGTTACAATCGTGCTACTAGCTACAACCAGAAGTTCAAAGGCAAGGC<br>CATATTTACTGTAGACAAATCCTCCAGCACAGCCTACATGCAATTCAACAGCCTGACATCTGAGGACTC<br>TGCTGTCTATTACTGTGCAAGA |
| IGHV-5LWB | IGHV2-6-5*01 | | F | CAGGTGCAGCTGAAGGAGTCAGGACCTGGCCTGGTGGCGCCCTCACAGAGCCTGTCCATCACATGCA<br>CTGTCTCAGGGTTCTCATTAACCGACTATGGTGTAAGCTGGATTCGCCAGCCTCCAGGAAAGGGTCTG<br>GAGTGGCTGGGAGTAATATGGGGTGGTGGAAGCACATACTATAATTCAGCTCTCAAATCCAGACTGA<br>GCATCAGCAAGGACAACTCCAAGAGCCAAGTTTTCTTAAAAATGAACAGTCTGCAAACTGATGACACA<br>GCCATGTACTACTGTGCCAAACA |
| IGHV-5O75 | | JACKSON: balbIGHV010 | F | CAGGTCCAGTTGCAGCAGTCTGGAGCTGAGCTGGTAAGGCCTGGGACTTCAGTGAAGATATCCTGCA<br>AGGCTTCTGGATACGCCTTCACTAACTACTGGCTAGGTTGGGTAAAGCAGAGGCCTGGACATGGACTT<br>GAGTGGATTGGAGATATTTACCCTGGAAGTGGTAATACTTACTACAATGAGAAGTTCAAGGGCAAAG<br>CCACACTGACTGCAGACAAATCCTCGAGCACAGCCTATATGCAGCTCAGTAGCCTGACATCTGAGGAC<br>TCTGCTGTCTATTTCTGTGCAAGA |
| IGHV-5UNX | | Haines: J558.34 | F | CAGGTCCAACTGCAGCAGCCTGGGGCTGAGCTGGTGAGGCCTGGGGTTTCAGTGAAGCTGTCCTGCA<br>AGGCTTCTGGCTACACATTCACCAGCTACTGGATGCACTGGATTAAGCAGAGGCCTGAGCAAGGCCTT<br>GAGAGGATTGGAGAGATTAATCCTAGCAATGGTGGTACTAACTACAATGAGAAGTTCAAGAGCAAGG<br>CCACACTGACTGTAGACAAATCCTCCAGCACAGCCTACATGCAACTCAGCAGCCTGACATCTGAGGAC<br>TCTGCGGTCTATTACTGTGCAAGA |

| | | | | |
|---|---|---|---|---|
| IGHV-5VOM | IGHV4-2*02 | | F | GAGGTGAAGCTTCTCGAGTCTGGAGGTGGCCTGGTGCAGCCTGGAGGATCCCTGAATCTCTCCTGTGC AGCCTCAGGATTCGATTTTAGTAGATACTGGATGAGTTGGGCTCGGCAGGCTCCAGGGAAAGGGCAG GAATGGATTGGAGAAATTAATCCAGGAAGCAGTACGATAAACTATACGCCATCTCTAAAGGATAAATT CATCATCTCCAGAGACAACGCCAAAAATACGCTGTACCTGCAAATGAGCAAAGTGAGATCTGAGGACA CAGCCCTTTATTACTGTGCAAGACT |
| IGHV-6D76 | IGHV14-1*02 | | F | GAGGTTCAGCTGCAGCAGTCTGGGGCTGAGCTTGTGAGGCCAGGGGCCTTAGTCAAGTTGTCCTGCA AAGCTTCTGGCTTCAACATTAAAGACTACTATATGCACTGGGTGAAGCAGAGGCCTGAACAGGGCCTG GAGTGGATTGGATGGATTGATCCTGAGAATGGTAATACTATATATGACCCGAAGTTCCAGGGCAAGG CCAGTATAACAGCAGACACATCCTCCAACACAGCCTACCTGCAGCTCAGCAGCCTGACATCTGAGGAC ACTGCCGTCTATTACTGTGCTAGA |
| IGHV-6NFG | | VBASE2: musIGHV023 | F | GAGGTCCAGCTGCAGCAGTCTGGACCTGAGCTGGTAAAGCCTGGGGCTTCAGTGAAGATGTCCTGCA AGGCTTCTGGATACACATTCACTAGCTATGTTATGCACTGGGTGAAGCAGAAGCCTGGGCAGGGCCTT GAGTGGATTGGATATATTAATCCTTACAATGATGGTACTAAGTACAATGAGAAGTTCAAAGGCAAGGC CACACTGACTTCAGACAAATCCTCCAGCACAGCCTACATGGAGCTCAGCAGCCTGACCTCTGAGGACT CTGCGGTCTATTACTGTGCAAGA |
| IGHV-6YTP | IGHV2-6*02 | | F | CAGGTGCAGCTGAAGGAGTCAGGACCTGGCCTGGTGGCGCCCTCACAGAGCCTGTCCATCACATGCA CCGTCTCAGGGTTCTCATTAACTAGCTATGGTGTACACTGGGTTCGCCAGCCTCCAGGAAAGGGTCTG GAGTGGCTGGTAGTGATATGGAGTGATGGAAGCACAACCTATAATTCAGCTCTCAAATCCAGACTGA GCATCAGCAAGGACAACTCCAAGAGCCAAGTTTTCTTAAAAATGAACAGTCTCCAAACTGATGACACA GCCATGTACTACTGTGCCAGAAA |
| IGHV-72VE | | JACKSON: balbIGHV011 | F | CAGGTCCAGCTGCAGCAGTCTGGACCTGAGCTGGTGAAGCCTGGGGCTTCAGTGAAGATATCCTGCA AGGCTTCTGGCTACAGCTTCACAAGCTACTATATACACTGGGTGAAGCAGAGGCCTGGACAGGGACTT GAGTGGATTGGATGGATTTTTCCTGGAAGTGGTAATACTAAGTACAATGAGAAGTTCAAGGGCAAGG CCACACTGACGGCAGACACATCCTCCAGCACAGCCTACATGCAGCTCAGCAGCCTGACATCTGAGGAC TCTGCAGTCTATTTCTGTGCAAGA |
| IGHV-7BBU | IGHV6-6*02 | | F | GAAGTGAAGCTTGAGGAGTCTGGAGGAGGCTTGGTGCAACCTGGAGGATCCATGAAACTCTCCTGTG TTGCCTCTGGATTCACTTTCAGTAACTACTGGATGAACTGGGTCCGCCAGTCTCCAGAGAAGGGGCTT GAGTGGGTTGCTGAAATTAGATTGAAATCTAATAATTATGCAACACATTATGCGGAGTCTGTGAAAGG GAGGTTCACCATCTCAAGAGATGATTCCAAAAGTAGTGTCTACCTGCAAATGAACAACTTAAGAGCTG AAGACACTGGCATTTATTACTGTACCAGG |
| IGHV-7MCI | IGHV7-3*02 | | F | GAGGTGAAGCTGGTGGAGTCTGGAGGAGGCTTGGTACAGCCTGGGGGTTCTCTGAGACTCTCCTGTG CAACTTCTGGGTTCACCTTCACTGATTACTACATGAGCTGGGTCCGCCAGCCTCCAGGAAAGGCACTTG AGTGGTTGGGTTTTATTAGAAACAAAGCTAATGGTTACACAACAGAGTACAGTGCATCTGTGAAGGGT |

| | | | | |
|---|---|---|---|---|
| | | | | CGGTTCACCATCTCCAGAGATAATTCCCAAAGCATCCTCTATCTTCAAATGAACACCCTGAGAGCTGAG GACAGTGCCACTTATTACTGTGCAAGAGATA |
| IGHV-7MU3 | IGHV1S122*01 | | F | CAGGTCCAACTCCAGCAGCCTGGGGCTGAACTGGTGAAGCCTGGGGCTTCAGTGAAGTTGTCCTGCA AGGCTTCTGGCTACACCTTCACCAGCTACTGGATGCACTGGGTGAAGCTGAGGCCTGGACAAGGCTTT GAGTGGATTGGAGAGATTAATCCTAGCAATGGTGGTACTAACTACAATGAGAAGTTCAAGAGAAAGG CCACACTGACTGTAGACAAATCCTCCAGCACAGCCTACATGCAACTCAGCAGCCTGACATCTGAGGAC TCTGCGGTCTATTACTGTACAATA |
| IGHV-7PG3 | | Haines: J558.20 | F | CAGGTCCAGCTGCAGCAGTCTGGAGATGATCTGGTAAAGCCTGGGGCCTCAGTGAAGCTGTCCTGCA AGGCTTCTGGCTACACCTTCACCAGCTACTGGATTAACTGGATAAAACAGAGGCCTGGACAGGGCCTT GAGTGGATAGGACGTATTGCTCCTGGAAGTGGTAGTACTTACTACAATGAAATGTTCAAGGGCAAGG CAACACTGACTGTAGACACATCCTCCAGCACAGCCTACATTCAGCTCAGCAGCCTGTCATCTGAGGACT CTGCTGTCTATTTCTGTGCAAGA |
| IGHV-7TJA | | JACKSON: balbIGHV003 | F | CAGGTTCAACTGCAGCAGTCTGGGGCTGAGCTGGTGAGGCCTGGGGCTTCAGTGAAGCTGTCCTGCA AGGCTTTGGGCTACACATTTACTGACTATGAAATGCACTGGGTGAAGCAGACACCTGTGCATGGCCTG GAATGGATTGGAGCTATTCATCCAGGAAGTGGTGGTACTGCCTACAATCAGAAGTTCAAGGGCAAGG CCACACTGACTGCAGACAAATCCTCCAGCACAGCCTACATGGAGCTCAGCAGCCTGACATCTGAGGAC TCTGCTGTCTATTACTGTACAAGA |
| IGHV-A3LZ | | Haines: J558.29 | F | CAGGTCCAGCTGCAGCAGTCTGGACCTGAGCTGGTGAAGCCTGGGGCTTCAGTGAGGATATCCTGCA AGGCTTCTGGCTACACCTTCACAAGCTACTATATACACTGGGTGAAGCAGAGGCCTGGACAGGGACTT GAGTGGATTGGATGGATTTATCCTGGAAATGTTAATACTAAGTACAATGAGAAGTTCAAGGGCAAGG CCACACTGACTGCAGACAAATCCTCCAGCACAGCCTACATGCAGCTCAGCAGCCTGACCTCTGAGGAC TCTGCGGTCTATTTCTGTGCAAGA |
| IGHV-AE2O | | JACKSON: balbIGHV012 | F | CAGGTCCAGTTGCAGCAGTCTGGCCTGAGCTGGTGAGGCCTGGGGTCTCAGTGAAGATTTCCTGCA AGGGTTCCAGCTACACATTCACTGATTATGCTATGCACTGGGTGAAGCAGAGTCATGCAAAGAGTCTA GAGTGGATTGGAGTTATTAGTACTTACTATGGTAATACTAACTACAACCAGAAGTTTAAGGGCAAGGC CACAATGACTGTAGACAAATCCTCCAGCACAGCCTATATGGAACTTGCCAGATTGACATCTGAGGATT CTGCCGTCTATTACTGTGCAAGA |
| IGHV-AICW | IGHV5-2*01 | | F | GAGGTGCAGCTGGTGGAGTCTGGGGGAGGCTTAGTGCAGCCTGGAGAGTCCCTGAAACTCTCCTGTG AATCCAATGAATACGAATTCCCTTCCCATGACATGTCTTGGGTCCGCAAGACTCCGGAGAAGAGGCTG GAGTTGGTCGCAGCCATTAATAGTGATGGTGGTAGCACCTACTATCCAGACACCATGGAGAGACGATT CATCATCTCCAGAGACAATACCAAGAAGACCCTGTACCTGCAAATGAGCAGTCTGAGGTCTGAGGACA CAGCCTTGTATTACTGTGCAAGACA |

| | | | | |
|---|---|---|---|---|
| IGHV-AK3H | | JACKSON: balbIGHV021 | F | GAAGTGAAGCTTGAGGAGTCTGGAGGAGGCTTGGTGCAACCTGGAGGATCCATGAAACTCTCCTGTG<br>TAGCCTCTGGATTTACTTTCAGTAGCTACTGGATGTCTTGGGTCCGCCAGTCTCCAGAGAAGGGGCTT<br>GAGTGGGTTGCTGAAATTAGATTGAAATCTGATAATTATGCAACACATTATGCGGAGTCTGTGAAAGG<br>GAAGTTCACCATCTCAAGAGATGATTCCAAAAGTCGTCTCTACCTGCAAATGAACAGCTTAAGAGCTG<br>AAGACACTGGAATTTATTACTGTACAGG |
| IGHV-APOU | IGHV1S34*01 | | F | GAGGTCCAGCTGCAGCAGTCTGGACCTGAGCTAGTGAAGACTGGGGCTTCAGTGAAGATATCCTGCA<br>AGGCTTCTGGTTACTCATTCACTGGTTACTACATGCACTGGGTCAAGCAGAGCCATGGAAAGAGCCTT<br>GAGTGGATTGGATATATTAGTTGTTACAATGGTGCTACTAGCTACAACCAGAAGTTCAAGGGCAAGGC<br>CACATTTACTGTAGACACATCCTCCAGCACAGCCTACATGCAGTTCAACAGCCTGACATCTGAAGACTC<br>TGCGGTCTATTACTGTGCAAGA |
| IGHV-AX4R | | JACKSON: balbIGHV038 | F | AAGGTCCAGCTGCAGCAGTCTGGAGCTGAGCTGGTGAAACCCGGGGCATCAGTGAAGCTGTCCTGCA<br>AGGCTTCTGGCTACACCTTCACTGAGTATATTATACACTGGGTAAAGCAGAGGTCTGGACAGGGTCTT<br>GAGTGGATTGGGTGGTTTTACCCTGGAAGTGGTAGTATAAAGTACAATGAGAAATTCAAGGACAAGG<br>CCACATTGACTGCGGACAAATCCTCCAGCACAGTCTATATGGAGCTTAGTAGATTGACATCTGAAGAC<br>TCTGCGGTCTATTTCTGTGCAAGACACGAAGA |
| IGHV-AX4W | | JACKSON: balbIGHV018 | F | CAGGTCCAACTGCAGCAGCCTGGGGCTGAGCTGGTGAAGCCTGGGGCTTCAGTGAAGATGTCCTGCA<br>AGGCTTCTGGCTACACCTTCACCAGCTACTGGATGCACTGGGTGAAGCAGAGGCCTGGACAAGGCCTT<br>GAGTGGATCGGAGTGATTGATCCTTCTGATAGTTATACTAGCTACAATCAAAAGTTCAAGGGCAAGGC<br>CACATTGACTGTAGACACATCCTCCAGCACAGCCTACATGCAGCTCAGCAGCCTGACATCTGAGGACT<br>CTGCGGTCTATTACTGTACAAGA |
| IGHV-AXRQ | | JACKSON: balbIGHV030 | F | CAGGTTACTCTGAAAGAGTCTGGCCCTGGGATATTGCAGCCCTCCCAGACCCTCAGTCTGACTTGTTCT<br>TTCTCTGGGTTTTCACTGAGCACTTCTGGTATGGGTGTAGGCTGGATTCGTCAGCCTTCAGGAGAGGG<br>TCTAGAGTGGCTGGCAGACATTTGGTGGGATGACAATAAGTACTATAACCCATCCCTGAAGAGCCGGC<br>TCACAATCTCCAAGGATACCTCCAGCAACCAGGTATTCCTCAAGATCACCAGTGTGGACACTGCAGATA<br>CTGCCACTTACTACTGTGCTCGAAGAG |
| IGHV-BDLN | IGHV5-6-2*01 | | F | GACGTGAAGCTCGTGGAGTCTGGGGGAGGCTTAGTGAAGCTTGGAGGGTCCCTGAAACTCTCCTGTG<br>CAGCCTCTGGATTCACTTTCAGTAGCTATTACATGTCTTGGGTTCGCCAGACTCCAGAGAAGAGGCTG<br>GAGTTGGTCGCAGCCATTAATAGTAATGGTGGTAGCACCTACTATCCAGACACTGTGAAGGGCCGATT<br>CACCATCTCCAGAGACAATGCCAAGAACACCCTGTACCTGCAAATGAGCAGTCTGAAGTCTGAGGACA<br>CAGCCTTGTATTACTGTGCAAGACA |
| IGHV-BIQU | | Haines: J558.32 | F | CAGGTTCAGCTGCAGCAGTCTGGAGCTGAACTGGTAAAGCCTGGGGCTTCAGTGAAGTTGTCCTGCA<br>AGGCTTCTGGCTACACCTTCACAAGCTATGATATAAACTGGGTGAGGCAGAGGCCTGAACAGGGACTT<br>GAGTGGATTGGATGGATTTTTCCTGGAGATGGTAGTACTAAGTACAATGAGAAGTTCAAGGGCAAGG |

| | | | | |
|---|---|---|---|---|
| | | | | CCACACTGACTACAGACAAATCCTCCAGCACAGCCTACATGCAGCTCAGCAGGCTGACATCTGAGGACTCTGCTGTCTATTTCTGTGCAAGA |
| IGHV-BISP | | VBASE2: musIGHV386 | F | CAGGTTCAGCTGCAGCAGTCTGGAGCTGAGCTGATGAAGCCTGGGGCCTCAGTGAAGATATCCTGCAAGGCTACTGGCTACACATTCAGTAGCTACTGGATAGAGTGGGTAAAGCAGAGGCCTGGACATGGCCTTGAGTGGATTGGAGAGATTTTACCTGGAAGTGGTAGTACTAACTACAATGAGAAGTTCAAGGGCAAGGCCACATTCACTGCAGATACATCCTCCAACACAGCCTACATGCAACTCAGCAGCCTGACATCTGAGGACTCTGCCGTCTATTACTGTGCAAGA |
| IGHV-BQCW | IGHV1-69*02 | | F | CAGGTCCAACTGCAGCAGCCTGGGGCTGAGCTTGTGAAGCCTGGGGCTTCAGTGAAGCTGTCCTGCAAGGCTTCTGGCTACACCTTCACCAGCTACTGGATGCACTGGGTGAAGCAGAGGCCTGGACAAGGCCTTGAGTGGATCGGAGAGATTGATCCTTCTGATAGTTATACTAACTACAATCAAAAGTTCAAGGGCAAGGCCACATTGACTGTAGACAAATCCTCCAGCACAGCCTACATGCAGCTCAGCAGCCTGACATCTGAGGACTCTGCGGTCTATTACTGTGCAAGA |
| IGHV-BT4I | IGHV1S10*01 | | P | CAGGTCCAGCTGCAGCAGTCTGGACCTGAGCTGGTGAGGCCTGGGACTTCAGTGAAGATATCCTGCAAGGCTTCTGGCTATACCTTCCTCACCTACTGGATGAACTGGGTGAAGTAGATGCCTGGACAGGGCCTTGAGTGGATTGGACAGATTTTTCCTGCAAGTGGTAGTACTAACTACAATGAGATGTTCAAGGGCAAGGCCACATTGACTGTAGACACATCCTCCAGCACAGCCTACATGCAGCTAAGCAGCCTGACATCTGAGGACTCTGCGGTCTATTTCTGTGCAAGA |
| IGHV-CFNH | | Haines: J558.6 | F | GAGGTCCTGCTGCAACAGTCTGGACCTGAGCTGGTGAAGCCTGGGGCTTCAGTGAAGATACCCTGCAAGGCTTCTGGATACACATTCACTGACTACAACATGGACTGGGTGAAGCAGAGCCATGGAAAGAGCCTTGAGTGGATTGGAGATATTAATCCTAACAATGGTGGTACTATCTACAACCAGAAGTTCAAGGGCAAGGCCACATTGACTGTAGACAAGTCCTCCAGCACAGCCTACATGGAGCTCCGCAGCCTGACATCTGAGGACACTGCAGTCTATTACTGTGCAAGA |
| IGHV-CSIM | IGHV12-3*02 | | F | CAGATGCAGCTTCAGGAGTCAGGACCTGGCCTGGTGAAACCCTCACAGTCACTCTTCCTCGCCTGCTCTATTACTGGTTTCCCCATCACCAGTGGTTACTACTGGATCTGGATCCGTCAGTCACCTGGGAAACCCCTAGAATGGATGGGGTACATCACTCATAGTGGGGAAACTTTCTACAACCCATCCCTCCAGAGCCCCATCTCCATTACTAGAGAAACATCCAAGAACCAGTTCTTTCTGCAATTGAACTCTGTGACCACAGAGGACACAGCCATGTATTACTGTGCAGGAGACAGA |
| IGHV-CWNQ | | JACKSON: balbIGHV008 | F | GAGGTTCAGCTCCAGCAGTCTGGGACTGTGCTGGCAAGGCCTGGGGCTTCCGTGAAGATGTCCTGCAAGGCTTCTGGCTACAGCTTTACCAGCTACTGGATGCACTGGGTAAAACAGAGGCCTGGACAGGGTCTAGAATGGATTGGTGCTATTTATCCTGGAAATAGTGATACTAGCTACAACCAGAAGTTCAAGGGCAAGGCCAAACTGACTGCAGTCACATCCGCCAGCACTGCCTACATGGAGCTCAGCAGCCTGACAAATGAGGACTCTGCGGTCTATTACTGTACAAGA |

| | | | | |
|---|---|---|---|---|
| IGHV-DMIT | | Haines: J558.52 | F | CAGGTTCAGCTGCAGCAGTCTGGGGCTGAGCTGGTGAAGCCTGGGGCCTCAGTGAAGATGTCCTGCA<br>AGGCTTTTGGCTACACCTTCACTACCTATCCAATAGAGTGGATGAAGCAGAATCATGGGAAGAGCCTA<br>GAGTGGATTGGAAATTTTCATCCTTACAATGATGATACTAAGTACAATGAAAAATTCAAGGGCAAGGC<br>CAAATTGACTGTAGAAAAATCCTCTAGCACAGTCTACTTGGAGCTCAGCCGATTAACATCTGATGACTC<br>TGCTGTTTATTACTGTGCAAGG |
| IGHV-DRA4 | | Haines: J558.22 | F | CAGGTTCAGCTGCAGCAGTCTGGACCTGAGCTGGTGAAGCCTGGGGCTTCAGTGAAGATGTCCTGCA<br>AGGCTTCTGGATACACATTCACTGACTATGTTATAAGCTGGGTGAAGCAGAGAACTGGACAGGGCCTT<br>GAGTGGATTGGAGAGATTTATCCTGGAAGTGGTAGTACTTACTACAATGAG AAGTTCAAGGGCAAGG<br>CCACACTGACTGCAGACAAATCCTCCAACACAGCCTACATGCAGCTCAGCAGCCTGACATCTGAGGAC<br>TCTGCGGTCTATTTCTGTGCAAGA |
| IGHV-DUV7 | IGHV14-2*02 | | P | TAGGTTAAGCTGCAGCAGTCTGGGGCAGAGCTTGTGAAGCCAGGGGCCTCAGTCAAGTTGTCCTGCA<br>AAGCTTCTGGCTTCAACATTAAAGACTACTATATGCACTGAGTGAAGCAGAGGCCTGAACAGGGCCTG<br>GAGTGGATTGGAAGGATTGATCCTGAGGATGGTGAAACTAAATATGCCCCGAAGTTCCAGGGCAAGG<br>CCACTATAACAGCAGACACATCCTCCAACACAGCCTACCTGCAGCTCAGCAGCCTGACATCTGAGGAC<br>ACTGCCGTCTATTACTGTGCTAGA |
| IGHV-EBWE | IGHV1S53*02 | | F | CAGGTTCAGCTGCAGCAGTCTGACGCTGAGTTGGTGAAACCTGGGGCTTCAGTGAAGATATCCTGCAA<br>GGCTTCTGGCTACACCTTCACTGACCATGCTATTCACTGGGTGAAGCAGAAGCCTGAACAGGGCCTGG<br>AATGGATTGGATATATTTCTCCCGGAAATGGTGATATTAAGTACAATGAGAAGTTCAAGGGCAAGGCC<br>ACACTGACTGCAGACAAATCCTCCAGCACTGCCTACATGCAGCTCAACAGCCTGACATCTGAGGATTCT<br>GCAGTGTATTTCTGTAAAAGA |
| IGHV-EK3O | IGHV1-84*02 | | F | CAGATCCAGCTGCAGCAGTCTGGACCTGAGCTGGTGAAGCCTGGGGCTTCAGTGAAGATATCCTGCA<br>AGGCTTCTGGCTACACCTTCACTGACTACTATATAAACTGGGTGAAGCAGAAGCCTGGACAGGGACTT<br>GAGTGGATTGGATGGATTTATCCTGGAAGCGGTAATACTAAGTACAATGAGAAGTTCAAGGGCAAGG<br>CCACATTGACTGTAGACACATCCTCCAGCACAGCCTACATGCAGCTCAGCAGCCTGACATCTGAGGAC<br>ACTGCTGTCTATTTCTGTGCAAGA |
| IGHV-ES4V | IGHV1S33*01 | | F | CAGGTTCAGCTGCAGCAGTCTGGACCTGAGCTGGTGAAGCCTGGGGCTTTAGTGAAGATATCCTGCA<br>AGGCTTCTGGTTACACCTTCACAAGCTACGATATAAACTGGGTGAAGCAGAGGCCTGGACAGGGACTT<br>GAGTGGATTGGATGGATTTATCCTGGAGATGGTAGTACTAAGTACAATGAGAAATTCAAGGGCAAGG<br>CCACACTGACTGCAGACAAATCCTCCAGCACAGCCTACATGCAGCTCAGCAGCCTGACTTCTGAGAACT<br>CTGCAGTCTATTTCTGTGCAAGA |
| IGHV-F3Z6 | IGHV9-4*02 | | F | CAGATCCAGTTGGTGCAGTCTGGACCTGAGCTGAAGAAGCCTGGAGAGACAGTCAGGATCTCCTGCA<br>AGGCTTCTGGGTATACCTTCACAACTGCTGGAATGCAGTGGGTGCAAAAGATGCCAGG AAAGGGTTT<br>GAAGTGGATTGGCTGGATAAACACCCACTCTGGAGTGCCAAAATATGCAGAAGACTTCAAGGGACGG |

| | | | | |
|---|---|---|---|---|
| | | | | TTTGCCTTCTCTTTGGAAACCTCTGCCAGCACTGCATATTTACAGATAAGCAACCTCAAAAATGAGGAC<br>ACGGCTACGTATTTCTGTGCGAGA |
| IGHV-F5K7 | | VBASE2: musIGHV707 | F | CAGGTCCAACTGCAGCAGCCTGGGGCTGAGCTTGTGATGCCTGGGGCTTCAGTGAAGATGTCCTGCA<br>AGGCTTCTGGCTACACATTCACTGACTACTGGATGCACTGGGTGAAGCAGAGGCCTGGACAAGGCCTT<br>GAGTGGATCGGAGCGATTGATACTTCTGATAGTTATACTAGCTACAATCAAAAGTTCAAGGGCAAGGC<br>CACATTGACTGTAGACGAATCCTCCAGCACAGCCTACATGCAGCTCAGCAGCCTGACATCTGAGGACT<br>CTGCGGTCTATTACTGTGCAAGA |
| IGHV-FM2S | IGHV2-4-1*01 | | F | CAGGTGCAGCTGAAGCAGTCAGGACCTGGCCTAGTGCAGCCCTCACAGAGCCTGTCCATCACCTGCAC<br>AGTCTCTGGTTTCTCATTAACTAGCTATGGTGTACACTGGGTTCGCCAGTCTCCAGGAAAGGGTCTGGA<br>GTGGCTGGGAGTGATATGGAGTGGTGGAAGCACAGACTATAATGCAGCTTTCATATCCAGACTGAGC<br>ATCAGCAAGGACAACTCCAAGAGCCAAGTTTTCTTTAAAATGAACAGTCTGCAAGCTGATGACACAGC<br>CATATACTACTGTGCCAGAAA |
| IGHV-FMN7 | IGHV3-6*02 | | F | GATGTACAGCTTCAGGAGTCAGGACCTGGCCTCGTGAAACCTTCTCAGTCTCTGTCTCTCACCTGCTCT<br>GTCACTGGCTACTCCATCACCAGTGGTTATTACTGGAACTGGATCCGGCAGTTTCCAGGAAACAAACT<br>GGAATGGATGGGCTACATAAGCTACGACGGTAGCAATAACTACAACCCATCTCTCAAAAATCGAATCT<br>CCATCACTCGTGACACATCTAAGAACCAGTTTTTCCTGAAGTTGAATTCTGTGACTACTGAGGACACAG<br>CTACATATTACTGTGCAAGAGA |
| IGHV-FNCD | IGHV5-6-3*01 | | F | GAGGTGCAGCTGGTGGAGTCTGGGGGAGGCTTAGTGCAGCCTGGAGGGTCCCTGAAACTCTCCTGTG<br>CAGCCTCTGGATTCACTTTCAGTAGCTATGGCATGTCTTGGGTTCGCCAGACTCCAGACAAGAGGCTG<br>GAGTTGGTCGCAACCATTAATAGTAATGGTGGTAGCACCTATTATCCAGACAGTGTGAAGGGCCGATT<br>CACCATCTCCAGAGACAATGCCAAGAACACCCTGTACCTGCAAATGAGCAGTCTGAAGTCTGAGGACA<br>CAGCCATGTATTACTGTGCAAGAGA |
| IGHV-FOTI | IGHV2-2*02 | | F | CAGGTGCAGCTGAAGCAGTCAGGACCTGGCCTAGTGCAGCCCTCACAGAGCCTGTCCATCACCTGCAC<br>AGTCTCTGGTTTCTCATTAACTAGCTATGGTGTACACTGGGTTCGCCAGTCTCCAGGAAAGGGTCTGGA<br>GTGGCTGGGAGTGATATGGAGTGGTGGAAGCACAGACTATAATGCAGCTTTCATATCCAGACTGAGC<br>ATCAGCAAGGACAATTCCAAGAGCCAAGTTTTCTTTAAAATGAACAGTCTGCAAGCTAATGACACAGC<br>CATATATTACTGTGCCAGAAA |
| IGHV-G6P5 | | JACKSON: balbIGHV009 | F | CAGGTCCAGCTGCAGCAGTCTGGAGCTGGGCTGGTGAAACCCGGGGCATCAGTGAAGCTGTCCTGCA<br>AGGCTTCTGGCTACACCTTCACTGAGTATATTATACACTGGGTAAAGCAGAGGTCTGGACAGGGTCTT<br>GAGTGGATTGGGTGGTTTTACCCTGGAAGTGGTAGTATAAAGTACAATGAGAAATTCAAGGACAAGG<br>CCACATTGACTGCGGACAAATCCTCCAGCACAGTCTATATGGAGCTTAGTAGATTGACATCTGAAGAC<br>TCTGCGGTCTATTTCTGTGCAAGACACGAAGA |

| | | | | |
|---|---|---|---|---|
| IGHV-GC2L | | VBASE2: musIGHV671 | F | GAGGTTCAGCTCCAGCAGTCTGGGACTGTGCTGGCAAGGCCTGGGGCTTCAGTGAAGATGTCCTGCA<br>AGGCTTCTGGCTACACCTTTACCAGCTACTGGATGCACTGGGTAAAACAGAGGCCTGGACAGGGTCTG<br>GAATGGATTGGCGCTATTTATCCTGGAAATAGTGATACTAGCTACAACCAGAAGTTCAAGGGCAAGGC<br>CAAACTGACTGCAGTCACATCCACCAGCACTGCCTACATGGAGCTCAGCAGCCTGACAAATGAGGACT<br>CTGCGGTCTATTACTGTACAAGA |
| IGHV-GFFP | | JACKSON: balbIGHV007 | F | GAGATCCAGCTGCAGCAGACTGGACCTGAGCTGGTGAAGCCTGGGGCTTCAGTGAAGATATCCTGCA<br>AGGCTTCTGGTTATTCATTCACTGACTACATCATGCTCTGGGTGAAGCAGAGCCATGGAAAGAGCCTT<br>GAGTGGATTGGAAATATTAATCCTTACTATGGTAGTACTAGCTACAATCTGAAGTTCAAGGGCAAGGC<br>CACATTGACTGTAGACAAATCTTCCAGCACAGCCTACATGCAGCTCAACAGTCTGACATCTGAGGACTC<br>TGCAGTCTATTACTGTGCAAGA |
| IGHV-GHLB | IGHV9-3-1*01 | | F | CAGATCCAGTTGGTGCAGTCTGGACCTGAGCTGAAGAAGCCTGGAGAGACAGTCAAGATCTCCTGCA<br>AGGCTTCTGGGTATACCTTCACAAACTATGGAATGAACTGGGTGAAGCAGGCTCCAGGAAAGGGTTT<br>AAAGTGGATGGGCTGGATAAACACCTACACTGGAGAGCCAACATATGCTGATGACTTCAAGGGACGG<br>TTTGCCTTCTCTTTGGAAACCTCTGCCAGCACTGCCTATTTGCAGATCAACAACCTCAAAAATGAGGAC<br>ACGGCTACATATTTCTGTGCAAGA |
| IGHV-GNPX | IGHV10-1*02 | | F | GAGGTGCAGCTTGTTGAGTCTGGTGGAGGATTGGTGCAGCCTAAAGGGTCATTGAAACTCTCATGTG<br>CAGCCTCTGGATTCACCTTCAATACCTACGCCATGAACTGGGTCCGCCAGGCTCCAGGAAAGGGTTTG<br>GAATGGGTTGCTCGCATAAGAAGTAAAAGTAATAATTATGCAACATATTATGCCGATTCAGTGAAAGA<br>CAGGTTCACCATCTCCAGAGATGATTCACAAAGCATGCTCTATCTGCAAATGAACAACTTGAAAACTGA<br>GGACACAGCCATGTATTACTGTGTGAGACA |
| IGHV-GPEO | | Haines: J558.13 | F | CAGGTCCAGCTGCAGCAGTCTGGAGCTGAGCTGGTAAGGCCTGGGACTTCAGTGAAGGTGTCCTGCA<br>AGGCTTCTGGATACGCCTTCACTAATTACTTGATAGAGTGGGTAAAGCAGAGGCCTGGACAGGGCCTT<br>GAGTGGATTGGAGTGATTAATCCTGGAAGTGGTGGTACTAACTACAATGAGAAGTTCAAGGGCAAGG<br>CAACACTGACTGCAGACAAATCCTCCAGCACTGCCTACATGCAGCTCAGCAGCCTGACATCTGATGACT<br>CTGCGGTTTATTTCTGTGCAAGC |
| IGHV-GRWZ | IGHV2-6-4*01 | | F | CAGGTGCAGCTGAAGGAGTCAGGACCTGGCCTGGTGGCACCCTCACAGAGCCTGTCCATCACATGCA<br>CTGTCTCTGGGTTCTCATTATCCAGATATAGTGTACACTGGGTTCGCCAGCCTCCAGGAAAGGGTCTGG<br>AGTGGCTGGGAATGATATGGGGTGGTGGAAGCACAGACTATAATTCAGCTCTCAAATCCAGACTGAG<br>CATCAGCAAGGACAACTCCAAGAGCCAAGTTTTCTTAAAAATGAACAGTCTGCAAACTGATGACACAG<br>CCATGTACTACTGTGCCAGAAA |
| IGHV-GSBQ | IGHV5-9-4*01 | | F | GAAGTGCAGCTGGTGGAGTCTGGGGGAGGCTTAGTGAAGCCTGGAGGGTCCCTGAAACTCTCCTGTG<br>CAGCCTCTGGATTCACTTTCAGTAGCTATGCCATGTCTTGGGTTCGCCAGTCTCCAGAGAAGAGGCTG<br>GAGTGGGTCGCAGAAATTAGTAGTGGTGGTAGTTACACCTACTATCCAGACACTGTGACGGGCCGATT |

| | | | | |
|---|---|---|---|---|
| | | | | CACCATCTCCAGAGACAATGCCAAGAACACCCTGTACCTGGAAATGAGCAGTCTGAGGTCTGAGGACA<br>CGGCCATGTATTACTGTGCAAGGGA |
| IGHV-GSYR | IGHV1-63*02 | | F | CAGGTCCAGCTGCAGCAGTCTGGAGCTGAGCTGGTAAGGCCTGGGACTTCAGTGAAGATGTCCTGCA<br>AGGCTGCTGGATACACCTTCACTAACTACTGGATAGGTTGGGTAAAGCAGAGGCCTGGACATGGCCTT<br>GAGTGGATTGGAGATATTTACCCTGGAGGTGGTTATACTAACTACAATGAGAAGTTCAAGGGCAAGG<br>CCACACTGACTGCAGACACATCCTCCAGCACAGCCTACATGCAGCTCAGCAGCCTGACATCTGAGGAC<br>TCTGCCATCTATTACTGTGCAAGA |
| IGHV-H2OA | IGHV7-1*02 | | F | GAGGTGAAGCTGGTGGAATCTGGAGGAGGCTTGGTACAGCCTGGGGGTTCTCTGAGACTCTCCTGTG<br>CAACTTCTGGGTTCACCTTCAGTGATTTCTACATGGAGTGGGTCCGCCAGCCTCCAGGGAAGAGACTG<br>GAGTGGATTGCTGCAAGTAGAAACAAAGCTAATGATTATACAACAGAGTACAGTGCATCTGTGAAGG<br>GTCGGTTCATCGTCTCCAGAGACACTTCCCAAAGCATCCTCTACCTTCAGATGAATGCCCTGAGAGCTG<br>AGGACACTGCCATTTATTACTGTGCAAGAGATGCA |
| IGHV-HHVI | | JACKSON: balbIGHV002 | F | CAGGTTCAACTGCAGCAGTCTGGGGCTGAGCTGGTGAGGCCTGGGGCTTCAGTGACGCTGTCCTGCA<br>AGGCTTCGGGCTACACATTTACTGACTATGAAATGCACTGGGTGAAGCAGACACCTGTGCATGGCCTG<br>GAATGGATTGGAGCTATTGATCCTGAAACTGGTGGTACTGCCTACAATCAGAAGTTCAAGGGCAAGG<br>CCACACTGACTGCAGACAAATCCTCCAGCACAGCCTACATGGAGCTCCGCAGCCTGACATCTGAGGAC<br>TCTGCCGTCTATTACTGTACAAGA |
| IGHV-HK55 | | VBASE2: musIGHV657 | F | CAGGTCCAGCTGCAGCAGTCTGGAGCTGAGCTGGTAAGGCCTGGGACTTCAGTGAAGGTGTCCTGCA<br>AGGCTTCTGGATACGCCTTCACTAATTACTTGATAGAGTGGGTAAAGCAGAGGCCTGGACAGGGCCTT<br>GAGTGGATTGGAGTGATTAATCCTGGAAGTGGTGGTACTAACTACAATGAGAAGTTCAAGGGCAAGG<br>CAACACTGACTGCAGACAAATCCTCCAGCACTGCCTACATGCAGCTCAGCAGCCTGACATCTGATGACT<br>CTGCGGTCTATTTCTGTGCAAGA |
| IGHV-HLLK | IGHV1-20*02 | | F | GAGGTTCAGCTGCAGCAGTCTGGACCTGAGCTGGTGAAGCCTGGGGCTTCAGTGAAGATATCCTGCA<br>AGGCTTCTGGTTACTCATTTACTGGCTACTTTATGAACTGGGTGATGCAGAGCCATGGAAAGAGCCTT<br>GAGTGGATTGGACGTATTAATCCTTACAATGGTGATACTTTCTACAACCAGAAGTTCAAGGGCAAGGC<br>CACATTGACTGTAGACAAATCCTCTAGCACAGCCCACATGGAGCTCCGGAGCCTGGCATCTGAGGACT<br>CTGCAGTCTATTATTGTGCAAGA |
| IGHV-HZPW | | VBASE2: musIGHV667 | F | CAGATTACTCTGAAAGAGTCTGGCCCTGGGATAGTGCAGCCATCCCAGCCCTTCAGACTTACTTGCACT<br>TTCTCTGGGTTTTCACTGAGCACTTCTGGTATAGGTGTAACCTGGATTCGTCAGCCCTCAGGGAAAGGT<br>CTGGAGTGGCTGGCAACGATTTGGTGGGATGATGATAACCGCTACAACCCATCTCTAAAGAGCAGGC<br>TCACAGTCTCCAAAGACACCTCCAACAACCAAGCATTCCTGAATATCATCACTGTGGAAACTGCAGATA<br>CTGCCATATACTACTGTGCTCAGAG |

| | | | | |
|---|---|---|---|---|
| IGHV-I463 | IGHV12-1-1*01 | | F | CAGATTCAGCTTAAGGAGTCTGGACCTGCTGTCATCAAGCCATCACAGTCACTGTCTCTCACCTGCATA GTCTCTGGATTCTCCATCACAAGTAGTAGTTATTGCTGGCACTGGATCCGCCAGCCCCCAGGAAAGGG GTTAGAGTGGATGGGGCGCATATGTTATGAAGGTTCAATATACTATAGTCCATCCATCAAAAGCCGCA GCACCATCTCCAGAGACACATCTCTGAACAAATTCTTTATCCAGCTGAGCTCTGTGACAAATGAGGACA CAGCCATGTACTACTGTTCCAGGGAAAACCA |
| IGHV-IAEK | IGHV2-3*01 | | F | CAGGTGCAGCTGAAGGAGTCAGGACCTGGCCTGGTGGCGCCCTCACAGAGCCTGTCCATCACATGCA CTGTCTCAGGGTTCTCATTAACCAGCTATGGTGTAAGCTGGGTTCGCCAGCCTCCAGGAAAGGGTCTG GAGTGGCTGGGAGTAATATGGGGTGACGGGAGCACAAATTATCATTCAGCTCTCATATCCAGACTGA GCATCAGCAAGGATAACTCCAAGAGCCAAGTTTTCTTAAAACTGAACAGTCTGCAAACTGATGACACA GCCACGTACTACTGTGCCAAACC |
| IGHV-ID7Q | | VBASE2: musIGHV668 | F | CAGGTCCAGCTGCAGCAGTCTGGGCCTGAGCTGGTGAGGCCTGGGGTCTCAGTGAAGATTTCCTGCA AGGGTTCCGGCTACACATTCACTGATTATGCTATGCACTGGGTGAAGCAGAGTCATGCAAAGAGTCTA GAGTGGATTGGAGTTATTAGTACTTACTATGGTAATACAAACTACAACCAGAAGTTTAAGGGCAAGGC CACAATGACTGTAGACAAATCCTCCAGCACAGCCTATATGGAACTTGCCAGATTGACATCTGAGGATT CTGCCATCTATTACTGTGCAAGA |
| IGHV-IMJE | IGHV2-5*01 | | F | CAGGTGCAGCTGAAGCAGTCAGGACCTGGCCTAGTGCAGCCCTCACAGAGCCTGTCCATAACCTGCAC AGTCTCTGGTTTCTCATTAACTAGCTATGGTGTACACTGGGTTCGCCAGTCTCCAGGAAAGGGTCTGGA GTGGCTGGGAGTGATATGGAGAGGTGGAAGCACAGACTACAATGCAGCTTTCATGTCCAGACTGAGC ATCACCAAGGACAACTCCAAGAGCCAAGTTTTCTTTAAAATGAACAGTCTGCAAGCTGATGACACTGC CATATACTACTGTGCCAAAAA |
| IGHV-IZSN | IGHV1S123*01 | | F | CAGGTCCAACTGCAGCAGCCTGGGGCTGAACTGGTGAAGCCTGGGGCTTCAGTGAAGTTGTCCTGCA AGGCTTCTGGCTACACCTTCACCAGCTACTATATGTACTGGGTGAAGCAGAGGCCTGGACAAGGCCTT GAGTGGATTGGGGGGATTAATCCTAGCAATGGTGGTACTAACTTCAATGAGAAGTTCAAGAGCAAGG CCACACTGACTGTAGACAAATCCTCCAGCACAGCCTACATGCAACTCAGCAGCCTGACATCTGAGGAC TCTGCGGTCTATTACTGTACAAGA |
| IGHV-JACH | IGHV6-7*02 | | F | GAGGTGAAGCTGGATGAGACTGGAGGAGGCTTGGTGCAACCTGGGAGGCCCATGAAACTCTCCTGT GTTGCCTCTGGATTCACTTTTAGTGACTACTGGATGAACTGGGTCCGCCAGTCTCCAGAGAAAGGACT GGAGTGGGTAGCACAAATTAGAAACAAACCTTATAATTATGAAACATATTATTCAGATTCTGTGAAAG GCAGATTCACCATCTCAAGAGATGATTCCAAAAGTAGTGTCTACCTGCAAATGAACAACTTAAGAGCT GAAGACATGGGTATCTATTACTGTACATGG |
| IGHV-JAPR | | JACKSON: balbIGHV006 | F | GAGGTTCAGCTGCAGCAGTCTGGACCTGAGCTGGTGAAGCCTGGGGCTTCAGTGAAGATATCCTGCA AGGCTTCTGGTTACTCATTTACTGGCTACTTTATGAACTGGGTGAAGCAGAGCCATGGAAAGAGCCTT GAGTGGATTGGACGTATTAATCCTTACAATGGTGATACTTTCTACAACCAGAAGTTCAAGGGCAAGGC |

| | | | | |
|---|---|---|---|---|
| | | | | CACATTGACTGTAGACAAATCCTCTAGCACAGCCCACATGGAGCTCCTGAGCCTGACATCTGAGGACTCTGCAGTCTATTATTGTGGAAGA |
| IGHV-JDDE | | JACKSON: balbIGHV023 | F | CAGGTTACTCTGAAAGAGTCTGGCCCTGGGATATTGCAGCCCTCCCAGACCCTCAGTCTGACTTGTTCTTTCTCTGGGTTTTCACTGAGCACTTCTGGTATGGGTGTGAGCTGGATTCGTCAGCCTTCAGGAAAGGGTCTGGAGTGGCTGGCACACATTTACTGGGATGATGACAAGCGCTATAACCCATCCCTGAAGAGCCGGCTCACAATCTCCAAGGATACCTCCAGAAACCAGGTATTCCTCAAGATCACCAGTGTGGACACTGCAGATACTGCCACATACTACTGTGCTCGAAGAG |
| IGHV-JDYC | IGHV2-2-2*01 | | F | CAGGTGCAGATGAAGCAGTCAGGACCTGGCCTAGTGCAGCCCTCACAGAGCCTGTCCATCACCTGCACAGTCTCTGGTTTCTCATTAACTAGCTATGGTGTACACTGGGTTCGCCAGTCTCCAGGAAAGGGTCTGGAGTGGCTGGGAGTGATATGGAGTGGTGGAAGCACAGACTATAATGCAGCTTTCATATCCAGACTGAGCATCAGCAAGGACAATTCCAAGAGCCAAGTTTTCTTTAAAATGAACAGTCTGCAAGCTGATGACACAGCCATATACTACTGTGTCAGAAA |
| IGHV-JHQJ | IGHV1-4*02 | | F | CAGGTCCAGCTGCAGCAGTCTGCAGCTGAACTGGCAAGACCTGGGGCCTCAGTGAAGATGTCCTGCAAGGCTTCTGGCTACACCTTTACTAGCTACACGATGCACTGGGTAAAACAGAGGCCTGGACAGGGTCTGGAATGGATTGGATACATTAATCCTAGCAGTGGATATACTGAGTACAATCAGAAGTTCAAGGACAAGACCACATTGACTGCAGACAAATCCTCCAGCACAGCCTACATGCAACTGAGCAGCCTGACATCTGAGGACTCTGCGGTCTATTACTGTGCAAGA |
| IGHV-JHWS | IGHV1-54*03 | | F | CAGGTCCAGTTGCAGCAGTCTGGAGCTGAGCTGGTAAGGCCTGGGACTTCAGTGAAGGTGTCCTGCAAGGCTTCTGGATACGCCTTCACTAATTACTTGATAGAGTGGGTAAAGCAGAGGCCTGGACAGGGCCTTGAGTGGATTGGGGTGATTAATCCTGGAAGTGGTGGTACTAACTACAATGAGAAGTTCAAGGGCAAGGCAACACTGACTGCAGACAAATCCTCCAGCACTGCCTACATGCAGCTCAGCAGCCTGACATCTGATGACTCTGCGGTCTATTTCTGTGCAAGA |
| IGHV-JRQS | IGHV5-4*02 | | F | GAAGTGCAGCTGGTGGAGTCTGGGGGAGGCTTAGTGAAGCCTGGAGGGTCCCTGAAACTCTCCTGTGCAGCCTCTGGATTCACTTTCAGTGACTATTACATGTATTGGGTTCGCCAGACTCCGGAAAAGAGGCTGGAGTGGGTCGCAACCATTAGTGATGGTGGTAGTTACACCTACTATCCAGACAGTGTGAAGGGGCGATTCACCATCTCCAGAGACAATGCCAAGAACAACCTGTACCTGCAAATGAGCAGTCTGAAGTCTGAGGACACAGCCATGTATTACTGTGCAAGAGA |
| IGHV-KO4L | IGHV1S22*01 | | F | CAGGTCCAACTGCAGCAACCTGGGTCTGAGCTGGTGAGGCCTGGAGCTTCAGTGAAGCTGTCCTGCAAGGCTTCTGGCTACACATTCACCAGCTACTGGATGCACTGGGTGAAGCAGAGGCATGGACAAGGCCTTGAGTGGATTGGAAATATTTATCCTGGTAGTGGTAGTACTAACTACGATGAGAAGTTCAAGAGCAAGGGCACACTGACTGTAGACACATCCTCCAGCACAGCCTACATGCACCTCAGCAGCCTGACATCTGAGGACTCTGCGGTCTATTACTGTACAAGA |

| | | | | |
|---|---|---|---|---|
| IGHV-KPEP | | VBASE2: musIGHV021 | F | CAGGTTCAGCTCCAGCAGTCTGGGGCTGAGCTGGCAAGACCTGGGGCTTCAGTGAAGTTGTCCTGCA<br>AGGCTTCTGGCTACACCTTTACTAGCTACTGGATGCAGTGGGTAAAACAGAGGCCTGGACAGGGTCTG<br>GAATGGATTGGGGCTATTTATCCTGGAGATGGTGATACTAGGTACACTCAGAAGTTCAAGGGCAAGG<br>CCACATTGACTGCAGATAAATCCTCCAGCACAGCCTACATGCAACTCAGCAGCTTGGCATCTGAGGAC<br>TCTGCGGTCTATTACTGTGCAAGA |
| IGHV-KQPM | | VBASE2: musIGHV578 | F | CAGGTGCAACTGCAGCAGTCTGGGCCTCAGCTGGTTAGGCCTGGGGCTTCAGTGAAGATATCCTGCA<br>AGGCTTCTGGTTACTCATTCACCAGCTACTGGATGCACTGGGTGAAGCAGAGGCCTGGACAAGGTCTT<br>GAGTGGATTGGCATGATTGATCCTTCCGATAGTGAAACTAGGTTAAATCAGAAGTTCAAGGACAAGG<br>CCACATTGACTGTAGACAAATCCTCCAGCACAGCCTACATGCAACTCAGCAGCCCGACATCTGAGGAC<br>TCTGCGGTCTATTACTGTGCAAGA |
| IGHV-KRG4 | | VBASE2: musIGHV396 | F | GAGGTCCAGCTGCAGCAGTCTGGACCTGAGCTGGAGAAGCCTGGCGCTTCAGTGAAGATATCCTGCA<br>AGGCTTCTGGTTACTCATTCACTGGCTACAACATGAACTGGGTGAAGCAGAGCAATGGAAAGAGCCTT<br>GAGTGGATTGGAAATATTGATCCTTACTATGGTGGTACTAGCTACAACCAGAAGTTCAAGGGCAAGGC<br>CACATTGACTGTAGACAAATCCTCCAGCACAGCCTACATGCAGCTCAAGAGCCTGACATCTGAGGACT<br>CTGCAGTCTATTACTGTGCAAGA |
| IGHV-KRVC | | JACKSON: balbIGHV037 | F | CAAGTTACTCTAAAAGAGTCTGGCCCTGGGATATTGAAGCCCTCACAGACCCTCAGTCTGACTTGTTCT<br>TTCTCTGGGTTTTCACTGAGCACTTCTGGTATGGGTGTAGGCTGGATTCGTCAGCCTTCAGGGAAGGG<br>TCTGGAGTGGCTGGCACACATTTGGTGGGATGATGATAAGTACTATAACCCATCCCTGAAGAGCCAGC<br>TCACAATCTCCAAGGATACCTCCAGAAACCAGGTATTCCTCAAGATCACCAGTGTGGACACTGCAGAT<br>ACTGCCACTTACTACTGTGCTCGAAGAG |
| IGHV-LAH4 | | VBASE2: musIGHV710 | F | CAGGTCCAACTGCAGCAGCCTGGGTCTGTGCTGGTGAGGCCTGGAGCTTCAGTGAAGCTGTCCTGCA<br>AGGCTTCTGGCTACACCTTCACCAGCTCCTGGATGCACTGGGCGAAGCAGAGGCCTGGACAAGGCCTT<br>GAGTGGATTGGAGAGATTCATCCTAATAGTGGTAATACTAACTACAATGAGAAGTTCAAGGGCAAGG<br>CCACACTGACTGTAGACACATCCTCCAGCACAGCCTACGTGGATCTCAGCAGCCTGACATCTGAGGAC<br>TCTGCGGTCTATTACTGTGCAAGA |
| IGHV-LLOE | IGHV4-1*02 | | F | GAGGTGAAGCTTCTCGAGTCTGGAGGTGGCCTGGTGCAGCCTGGAGGATCCCTGAAACTCTCCTGTG<br>CAGCCTCAGGATTCGATTTTAGTAGATACTGGATGAGTTGGGTCCGGCAGGCTCCAGGGAAAGGGCT<br>AGAATGGATTGGAGAAATTAATCCAGATAGCAGTACGATAAACTATACGCCATCTCTAAAGGATAAAT<br>TCATCATCTCCAGAGACAACGCCAAAAATACGCTGTACCTGCAAATGAGCAAAGTGAGATCTGAGGAC<br>ACAGCCCTTTATTACTGTGCAAGACC |
| IGHV-LP2Z | IGHV1S121*01 | | F | CAGGTGCAACTGCAGCAGCCTGGGGCTGAGCTGGTGAAGCCTGGGGCCTCAGTGAAGATGTCCTGCA<br>AGGCTTCTGGCTACACATTTACCAGTTACAATATGCACTGGGTAAAGCAGACACCTGGACAGGGCCTG<br>GAATGGATTGGAGCTATTTATCCAGGAAATGGTGATACTTCCTACAATCAGAAGTTCAAAGGCAAGGC |

| | | | | |
|---|---|---|---|---|
| | | | | CACATTGACTGCAGACAAATCCTCCAGCACAGCCTACATGCAGCTCAGCAGCCTGACATCTGAGGACT<br>CTGCGGTCTATTACTGTGCAAGA |
| IGHV-LPGY | IGHV5-9-1*01 | | F | GAAGTGATGCTGGTGGAGTCTGGGGGAGGCTTAGTGAAGCCTGGAGGGTCCCTGAAACTCTCCTGTG<br>CAGCCTCTGGATTCACTTTCAGTAGCTATGCCATGTCTTGGGTTCGCCAGACTCCGGAGAAGAGGCTG<br>GAGTGGGTCGCAACCATTAGTAGTGGTGGTAGTTACACCTACTATCCAGACAGTGTGAAGGGGCGAT<br>TCACCATCTCCAGAGACAATGCCAAGAACACCCTGTACCTGCAAATGAGCAGTCTGAGGTCTGAGGAC<br>ACGGCCATGTATTACTGTGCAAGA |
| IGHV-MCP4 | IGHV5-9*03 | | F | GAAGTGATGCTGGTGGAGTCTGGGGGAGGCTTAGTGAAGCCTGGAGGGTCCCTGAAACTCTCCTGTG<br>CAGCCTCTGGATTCACTTTCAGTAGCTATACCATGTCTTGGGTTCGCCAGACTCCGGAGAAGAGGCTG<br>GAGTGGGTCGCAACCATTAGTAGTGGTGGTGGTAACACCTACTATCCAGACAGTGTGAAGGGTCGAT<br>TCACCATCTCCAGAGACAATGCCAAGAACAACCTGTACCTGCAAATGAGCAGTCTGAGGTCTGAGGAC<br>ACGGCCTTGTATTACTGTGCAAGATA |
| IGHV-MG5V | IGHV13-2*02 | | F | CAGGTGCAGCTTGTAGAGACCGGGGGAGGCTTGGTGAGGCCTGGAAATTCTCTGAAACTCTCCTGTG<br>TTACCTCGGGATTCACTTTCAGTAACTACCGGATGCACTGGCTTCGCCAGCCTCCAGGGAAGAGGCTG<br>GAGTGGATTGCTGTAATTACAGTCAAATCTGATAATTATGGAGCAAATTATGCAGAGTCTGTGAAAGG<br>CAGATTCACTATTTCAAGAGATGATTCAAAAAGCAGTGTCTACCTGCAGATGAACAGATTAAGAGAGG<br>AAGACACTGCCACTTATTATTGTAGTAGA |
| IGHV-MGBE | | JACKSON: balbIGHV001 | F | GAGGTGCAGCTTGTTGAGTCTGGTGGAGGATTGGTGCAGCCTAAAGGATCATTGAAACTCTCATGTGC<br>CGCCTCTGGTTTCACCTTCAATACCTATGCCATGCACTGGGTCTGCCAGGCTCCAGGAAAGGGTTTGGA<br>ATGGGTTGCTCGCATAAGAAGTAAAAGTAATAATTATGCAACATATTATGCCGATTCAGTGAAAGACA<br>GATTCACCATCTCCAGAGATGATTCACAAAGCATGCTCTATCTGCAAATGAACAACCTGAAAACTGAG<br>GACACAGCCATGTATTACTGTGTGAGAGA |
| IGHV-ML5T | IGHV9-2-1*01 | | F | CAGATCCAGTTGGTGCAGTCTGGACCTGAGCTGAAGAAGCCTGGAGAGACAGTCAAGATCTCCTGCA<br>AGGCTTCTGGTTATACCTTCACAGACTATTCAATGCACTGGGTGAAGCAGGCTCCAGGAAAGGGTTTA<br>AAGTGGATGGGCTGGATAAACACTGAGACTGGTGAGCCAACATATGCAGATGACTTCAAGGGACGGT<br>TTGCCTTCTCTTTGGAAACCTCTGCCAGCACTGCCTATTTGCAGATCAACAACCTCAAAAATGAGGACA<br>CGGCTACATATTTCTGTGCTAGA |
| IGHV-NNFU | IGHV5-9-3*01 | | F | GAAGTGCAGCTGGTGGAGTCTGGGGGAGGCTTAGTGAAGCCTGGAGGGTCCCTGAAACTCTCCTGTG<br>CAGCCTCTGGATTCACTTTCAGTAGCTATGCCATGTCTTGGGTTCGCCAGACTCCGGAGAAGAGGCTG<br>GAGTGGGTCGCAACCATTAGTAGTGGTGGTAGTTACACCTACTATCCAGACAGTGTGAAGGGTCGATT<br>CACCATCTCCAGAGACAATGCCAAGAACACCCTGTACCTGCAAATGAGCAGTCTGAGGTCTGAGGACA<br>CGGCCATGTATTACTGTGCAAGACA |

| | | | | |
|---|---|---|---|---|
| IGHV-NPBL | | VBASE2: musIGHV592 | F | CAGGTCCAGCTGAAGCAGTCTGGAGCTGAGCTGGTGAGGCCTGGGGCTTCAGTGAAGCTGTCCTGCA<br>AGACTTCTGGATACATCTTCACCAGCTACTGGATTCACTGGGTAAAACAGAGGTCTGGACAGGGCCTT<br>GAGTGGATTGCAAGGATTTATCCTGGAACTGGTAGTACTTACTACAATGAGAAGTTCAAGGGCAAGG<br>CCACACTGACTGCAGACAAATCCTCCAGCACTGCCTACATGCAGCTCAGCAGCCTGAAATCTGAGGAC<br>TCTGCTGTCTATTTCTGTGCAAGA |
| IGHV-NUDM | | JACKSON: balbIGHV041 | OR<br>F | CAGGTTCAGCTCCAGCAGTCTGGGCCTGAGCTGGCAAGGCCTTGGGCTTCAGTGAAGATATCCTGCCA<br>GGCTTTCTACACCTTTTCCAGAAGGGTGTACTTTGCCATTAGGGATACCAACTACTGGATGCAGTGGGT<br>AAAACAGAGGCCTGGACAGGGTCTGGAATGGATCGGGGCTATTTATCCTGGAAATGGTGATACTAGT<br>TACAATCAGAAGTTCAAGGGCAAGGCCACATTGACTGCAGACAAATCCTCCAGCACAGCCTACATGCA<br>ACTCAGCAGCCTGACATCTGAGGACTCTGCGGTCTATTACTGTGCATGA |
| IGHV-NY4F | IGHV3-8*02 | | F | GAGGTGCAGCTTCAGGAGTCAGGACCTAGCCTCGTGAAACCTTCTCAGACTCTGTCCCTCACCTGTTCT<br>GTCACTGGCGACTCCATCACCAGTGGTTACTGGAACTGGATCCGGAAATTCCCAGGGAATAAACTTGA<br>GTACATGGGGTACATAAGCTACAGTGGTAGCACTTACTACAATCCATCTCTCAAAAGTCGAATCTCCAT<br>CACTCGAGACACATCCAAGAACCAGTACTACCTGCAGTTGAATTCTGTGACTACTGAGGACACAGCCA<br>CATATTACTGTGCAAGATA |
| IGHV-O4YA | IGHV3-2*02 | | F | GATGTGCAGCTTCAGGAGTCGGGACCTGGCCTGGTGAAACCTTCTCAGTCTCTGTCCCTCACCTGCACT<br>GTCACTGGCTACTCAATCACCAGTGATTATGCCTGGAACTGGATCCGGCAGTTTCCAGGAAACAAACT<br>GGAGTGGATGGGCTACATAAGCTACAGTGGTAGCACTAGCTACAACCCATCTCTCAAAAGTCGAATCT<br>CTATCACTCGAGACACATCCAAGAACCAGTTCTTCCTGCAGTTGAATTCTGTGACTACTGAGGACACAG<br>CCACATATTACTGTGCAAGA |
| IGHV-OCOS | | JACKSON: balbIGHV013 | F | CAGGTCCAGCTGCAGCAGTCTGGGCCTGAGGTGGTGAGGCCTGGGGTCTCAGTGAAGATTTCCTGCA<br>AGGGTTCCGGCTACACATTCACTGATTATGCTATGCACTGGGTGAAGCAGAGTCATGCAAAGAGTCTA<br>GAGTGGATTGGAGTTATTAGTACTTACAATGGTAATACAAACTACAACCAGAAGTTTAAGGGCAAGGC<br>CACAATGACTGTAGACAAATCCTCCAGCACAGCCTATATGGAACTTGCCAGATTGACATCTGAGGATT<br>CTGCCATCTATTACTGTGCAAGA |
| IGHV-OK63 | | VBASE2: musIGHV702 | F | CAGGTCCAACTGCAGCAACCTGGGTCTGAGCTGGTGAGGCCTGGAGCTTCAGTGAAGCTGTCCTGCA<br>AGGCTTCTGGCTACACATTCACCAGCTACTGGATGCACTGGGTGAAGCAGAGGCCTGGACAAGGCCTT<br>GAGTGGATTGGAAATATTTATCCTGGTAGTGGTAGTACTAACTACGATGAGAAGTTCAAGAGCAAGG<br>CCACACTGACTGTAGACACATCCTCCAGCACAGCCTACATGCAGCTCAGCAGCCTGACATCTGAGGAC<br>TCTGCGGTCTATTACTGTACAAGA |
| IGHV-OW3E | | VBASE2: musIGHV559 | F | CAGGTTCAGCTGCAGCAGTCTGGGGCTGAGCTGGTGAGGCCTGGGTCCTCAGTGAAGATTTCCTGCA<br>AGGCTTCTGGCTATGCATTCAGTAGCTACTGGATGAACTGGGTGAAGCAGAGGCCTGGACAGGGTCT<br>TGAGTGGATTGGACAGATTTATCCTGGAGATGGTGATACTAACTACAATGGAAAGTTCAAGGGTAAA |

| | | | | |
|---|---|---|---|---|
| | | | | GCCACACTGACTGCAGACAAATCCTCCAGCACAGCCTACATGCAGCTCAGCAGCCTAACATCTGAGGACTCTGCGGTCTATTTCTGTGCAAGA |
| IGHV-P2CQ | IGHV1S126*01 | | F | CAGGTGCAACTGCAGCAGCCTGGGGCTGAGCTGGTGAAGCCTGGGGCTTCAGTGAAGATATCCTGCAAGGCTTCTGGCTACACCTTCACCAGCTACTGGATGAACTGGGTGAAGCAGAGGCCTGGACAAGGCCTTGAGTGGATCGGAGAGATCGATCCTTCTGATAGTTATACTAACAACAATCAAAAGTTCAAGGACAAGGCCACATTGACTGTAGACAAATCCTCCAGCACAGCCTACATGCAGCTCAGCAGCCTGACATCTGAGGACTCTGCGGTCTATTACTGTGCAAGA |
| IGHV-P7AN | IGHV2-6-1*01 | | F | CAGGTGCAGCTGAAGGAGTCAGGACCTGGCCTGGTGGCGCCCTCACAGAGCCTGTCCATCACATGCACCATCTCAGGGTTCTCATTAACCAGCTATGGTGTACACTGGGTTCGCCAGCCTCCAGGAAAGGGTCTGGAGTGGCTGGTAGTGATATGGAGTGATGGAAGCACAACCTATAATTCAGCTCTCAAATCCAGACTGAGCATCAGCAAGGACAACTCCAAGAGCCAAGTTTTCTTAAAAATGAACAGTCTCCAAACTGATGACACAGCCATGTACTACTGTGCCAGACA |
| IGHV-PACA | IGHV10S3*01 | | F | GAGGTGCAGCTTGTTGAGACTGGTGGAGGATTGGTGCAGCCTAAAGGGTCATTGAAACTCTCATGTGCAGCCTCTGGATTCACCTTCAATACCAATGCCATGAACTGGGTCCGCCAGGCTCCAGGAAAGGGTTTGGAATGGGTTGCTCGCATAAGAAGTAAAAGTAATAATTATGCAACATATTATGCCGATTCAGTGAAAGACAGGTTCACCATCTCCAGAGATGATTCACAAAGCATGCTCTATCTGCAAATGAACAACTTGAAAACTGAGGACACAGCCATGTATTACTGTGTGAGAGA |
| IGHV-PEXG | | VBASE2: musIGHV560 | F | CAGGTCCAGCTGCAGCAGTCTGGGGCTGAGCTGGTGAGGCCTGGGGTCTCAGTGAAGATTTCCTGCAAGGGTTCTGGCTACACATTCACTGATTATGCTATGCACTGGGTGAAGCAGAGTCATGCAAAGAGTCTAGAGTGGATTGGAGTTATTAGTACTTACTATGGTGATGCTAGCTACAACCAGAAGTTCAAGGGCAAGGCCACAATGACTGTAGACAAATCCTCCAGCACAGCCTATATGGAACTTGCCAGACTGACATCTGAGGATTCTGCCATCTATTACTGTGCAAAA |
| IGHV-PRC5 | IGHV9-1*02 | | F | CAGATCCAGTTGGTGCAGTCTGGACCTGAGCTGAAGAAGCCTGGAGAGACAGTCAAGATCTCCTGCAAGGCTTCTGGGTATACCTTCACAAACTATGGAATGAACTGGGTGAAGCAGGCTCCAGGAAAGGGTTTAAAGTGGATGGGCTGGATAAACACCTACACTGGAGAGCCAACATATGCTGATGACTTCAAGGGACGGTTTGCCTTCTCTTTGGAAACCTCTGCCAGCACTGCCTATTTGCAGATCAACAACCTCAAAAATGAGGACATGGCTACATATTTCTGTGCAAGA |
| IGHV-PRJX | | JACKSON: balbIGHV022 | F | GAAGTGAAGCTTGAGGAGTCTGGAGGAGGCTTGGTGCAACCTGGAGGATCCATGAAACTCTCTTGTGCTGCCTCTGGATTCACTTTTAGTGACGCCTGGATGGACTGGGTCCGCCAGTCTCCAGAGAAGGGGCTTGAGTGGGTTGCTGAAATTAGAAGCAAAGCTAATAATCATGCAACATACTATGCTGAGTCTGTGAAAGGGAGGTTCACCATCTCAAGAGATGATTCCAAAAGTAGTGTCTACCTGCAAATGAACAGCTTAAGAGCTGAAGACACTGGCATTTATTACTGTACCAGG |

| | | | | |
|---|---|---|---|---|
| IGHV-QC3C | IGHV2-9-2*01 | | F | CAGGTGCAACTGAAGGAGTCAGGACCTGGCCTGGTGGCGCCCTCACAGAGCCTGTCCATTACCTGCACTGTCTCTGGGTTCTCATTAACCAGCTATGATATAAGCTGGATTCGCCAGCCACCAGGAAAGGGTCTGGAGTGGCTTGGAGTAATATGGACTGGTGGAGGCACAAATTATAATTCAGCTTTCATGTCCAGACTGAGCATCAGCAAGGACAACTCCAAGAGCCAAGTTTTCTTAAAAATGAACAGTCTGCAAACTGATGACACAGCCATATATTACTGTGTAAGAGA |
| IGHV-QFGX | | JACKSON: balbIGHV034 | F | CAGGTTACTCTGAAAGAGTCTGGCCCTGGGATATTGCAGCCCTCCCAGACCCTCAGTCTGACTTGTTCTTTCTCTGGGTTTTCACTGAGCACTTCTGGTATGGGTGTAGGCTGGATTCGTCAGCCTTCAGGGAAGGGTCTGGAGTGGCTGGCACACATTTGGTGGGATGATGATAAGTACTATAACACAGCCCTGAAGAGCGGGCTCACAATCTCCAAGGATACCTCCAAAAACCAGGTCTTCCTCAAGATCGCCAGTGTGGACACTGCAGATACTGCCACATACTACTGTGCTCGAATAG |
| IGHV-QG3Z | | JACKSON: balbIGHV020 | F | GAGGTCCAGCTTCAGCAGTCAGGACCTGAGCTGGTGAAACCTGGGGCCTCAGTGAAGATATCCTGCAAGGCTTCTGGATACACATTCACTGACTACAACATGCACTGGGTGAAGCAGAGCCATGGAAAGAGCCTTGAGTGGATTGGATATATTTATCCTTACAATGGTGGTACTGGCTACAACCAGAAGTTCAAGAGCAAGGCCACATTGACTGTAGACAATTCCTCCAGCACAGCCTACATGGAGCTCCGCAGCCTGACATCTGAGGACTCTGCAGTCTATTACTGTGCAAGA |
| IGHV-QK6G | | JACKSON: balbIGHV024 | F | CAGGTTACTCTGAAAGTGTCTGGCCCTGGGATATTGCAGCCATCACAGACTCTCGGCCTGGCCTGTACTTTCTCTGGGATTTCACTGAGTACTTCTGGTATGGGTTTGAGCTGGCTTCGTAAGCCCTCAGGGAAGGCTTTAGAGTGGCTGGCAAGCATTTGGAATAATGATAACTACTACAACCCATCTTTGAAGAGCCGGCTCACAATCTCCAAGGAGACCTCCAACAACCAAGTATTCCTTAAACTCACCAGTGTGGACACTGCAGATTCTACCACATACTACTGTGCTTGGAGAGAG |
| IGHV-QL4G | IGHV5-9*02 | | F | GAAGTGAAGCTGGTGGAGTCTGGGGGAGGCTTAGTGAAGCCTGGAGGGTCCCTGAAACTCTCCTGTGCAGCCTCTGGATTCGCTTTCAGTAGCTATGACATGTCTTGGGTTCGCCAGACTCCGGAGAAGAGGCTGGAGTGGGTCGCAACCATTAGTAGTGGTGGTAGTTACACCTACTATCCAGACAGTGTGAAGGGCCGATTCACCATCTCCAGAGACAATGCCAGGAACACCCTGTACCTGCAAATGAGCAGTCTGAGGTCTGAGGACACGGCCTTGTATTACTGTGCAAGACA |
| IGHV-QODY | IGHV5-12*02 | | F | GAAGTGAAGCTGGTGGAGTCTGGGGGAGGCTTAGTGCAGCCTGGAGGGTCCCTGAAACTCTCCTGTGCAACCTCTGGATTCACTTTCAGTGACTATTACATGTATTGGGTTCGCCAGACTCCAGAGAAGAGGCTGGAGTGGGTCGCATACATTAGTAATGGTGGTGGTAGCACCTATTATCCAGACACTGTAAAGGGCCGATTCACCATCTCCAGAGACAATGCCAAGAACACCCTGTACCTGCAAATGAGCCGTCTGAAGTCTGAGGACACAGCCATGTATTACTGTGCAAGACA |
| IGHV-QQ46 | | VBASE2: musIGHV588 | F | CAGGTCCAACTGCAGCAGCCTGGGGCTGAGCTGGTGAGGCCTGGGGCTTCAGTGAAGCTGTCCTGCAAGGCTTCTGGCTACACCTTCACCAGCTACTGGATGAACTGGGTGAAGCAGAGGCCTGGACAAGGCCTTGAATGGATTGGTATGATTGATCCTTCAGACAGTGAAACTCACTACAATCAAATGTTCAAGGACAAGGC |

| | | | | |
|---|---|---|---|---|
| | | | | CACATTGACTGTAGACAAATCCTCCAGCACAGCCTACATGCAGCTCAGCAGCCTGACATCTGAGGACTCTGCGGTCTATTACTGTGCAAGA |
| IGHV-R4HE | | JACKSON: balbIGHV028 | F | CAGGTTACTCTGAAAGAGTCTGGCCCTGGGATATTGCAGCCCTCCCAGACCCTCAGTCTGACTTGTTCTTTCTCTGGGTTTTCACTGAGCACTTCTGGTATGAGTGTAGGCTGGATTCGTCAGCCTTCAGGGAAGGGTCTGGAGTGGCTGGCACACATTTGGTGGAATGATGATAAGTACTATAACCCAGCCCTGAAAAGCCGGCTCACAATCTCCAAGGATACCTCCAACAACCAGGTATTCCTCAAGATCGCCAGTGTGGTCACTGCAGATACTGCCACATACTACTGTGCTCGAATAG |
| IGHV-R7KA | | VBASE2: musIGHV480 | F | GAGGTCCAGCTGCAACAGTCTGGACCTGAGCTGGTGAAGCCTGGGGCTTCAGTGAAGATATCCTGCAAGGCTTCTGGTTACTCATTCACTGGCTACTACATGCACTGGGTGAAGCAAAGCCATGTAAAGAGCCTTGAGTGGATTGGACGTATTAATCCTTACAATGGTGCTACTAGCTACAACCAGAATTTCAAGGACAAGGCCAGCTTGACTGTAGATAAGTCCTCCAGCACAGCCTACATGGAGCTCCACAGCCTGACATCTGAGGACTCTGCAGTCTATTACTGTGCAAGA |
| IGHV-RKXL | IGHV5-6*01 | | F | GAGGTGCAGCTGGTGGAGTCTGGGGGAGACTTAGTGAAGCCTGGAGGGTCCCTGAAACTCTCCTGTGCAGCCTCTGGATTCACTTTCAGTAGCTATGGCATGTCTTGGGTTCGCCAGACTCCAGACAAGAGGCTGGAGTGGGTCGCAACCATTAGTAGTGGTGGTAGTTACACCTACTATCCAGACAGTGTGAAGGGGCGATTCACCATCTCCAGAGACAATGCCAAGAACACCCTGTACCTGCAAATGAGCAGTCTGAAGTCTGAGGACACAGCCATGTATTACTGTGCAAGACA |
| IGHV-RLDP | | VBASE2: musIGHV483 | F | CAGGTCCAACTGCAGCAGTCTGGGCCTGAGCTGGTGAGGCCTGGGGCTTCAGTGAAGATGTCCTGCAAGGCTTCAGGCTATACCTTCACCAGCTACTGGATGCACTGGGTGAAACAGAGGCCTGGACAAGGCCTTGAGTGGATTGGCATGATTGATCCTTCCAATAGTGAAACTAGGTTAAATCAGAAGTTCAAGGACAAGGCCACATTGAATGTAGACAAATCCTCCAACACAGCCTACATGCAGCTCAGCAGCCTGACATCTGAGGACTCTGCAGTCTATTACTGTGCAAGA |
| IGHV-RNDH | | VBASE2: musIGHV532 | F | CAGGTCCAGCTTCAGCAGTCTGGGGCTGAACTGGCAAAACCTGGGGCCTCAGTGAAGATGTCCTGCAAGGCTTCTGGCTACACCTTTACTAGCTACTGGATGCACTGGGTAAAACAGAGGCCTGGACAGGGTCTGGAATGGATTGGATACATTAATCCTAGCACTGGTTATACTGAGTACAATCAGAAGTTCAAGGACAAGGCCACATTGACTGCAGACAAATCCTCCAGCACAGCCTACATGCAACTGAGCAGCCTGACATCTGAGGACTCTGCAGTCTATTACTGTGCAAGA |
| IGHV-RNSW | IGHV2-2-1*01 | | P | CAGGTGCAGCTGAAGCAGTCAGGACCTGGCCTAGTGCAGCCCTCACAGAGCCTGTCCATCACCTGCACATTCTCTGGTTTCTGATTAACCAGCTATGGTGTACACTGGGAGCGCCATTCTCCAGGAAAGGGTCTGGAGTGGCTGGGAGTGATATGGAGTGGTGTACACACAGACTATAATGCAGCTTTCATATCCAGATTGAGCATCAGCAAGGACAACTCCAAGAGCCAAGTTTTCTTTAAAATGAACAGTCTGCAAGCTGATGACACAGCCATATACTACTGTGCCAGAAA |

| | | | | |
|---|---|---|---|---|
| IGHV-ROYG | IGHV1S113*01 | | F | GAGGTCCAGCTGCAACAGTCTGGACCTGAGCTGGTGAAGCCTGGGGCTTCAGTGAAGATATCCTGCA<br>AGACTTCTGGATACACATTCACTGAATACACCATGCACTGGGTGAAGCAGAGCCATGGAAAGAGCCTT<br>GAGTGGATTGGAGGTATTAATCCTAACAATGGTGGTACTAGCTACAACCAGAAGTTCAAGGGCAAGG<br>CCACATTGACTGTAGACAAGTCCTCCAGCACAGCCTACATGGAGCTCCGCAGCCTGACATCTGAGGAT<br>TCTGCAGTCTATTACTGTGCAAGA |
| IGHV-RRLD | IGHV5-12-2*01 | | F | GAAGTGAAGCTGGTGGAGTCTGGGGGAGGTTTAGTGCAGCCTGGAGGGTCCCTGAAACTCTCCTGTG<br>CAGCCTCTGGATTCACTTTCAGTAGCTATACCATGTCTTGGGTTCGCCAGACTCCAGAGAAGAGGCTG<br>GAGTGGGTCGCATACATTAGTAATGGTGGTGGTAGCACCTACTATCCAGACACTGTAAAGGGCCGATT<br>CACCATCTCCAGAGACAATGCCAAGAACACCCTGTACCTGCAAATGAGCAGTCTGAAGTCTGAGGACA<br>CGGCCATGTATTACTGTGCAAGACA |
| IGHV-RYF3 | | VBASE2: musIGHV433 | F | CAGGTTACTCTGAAAGAGTCTGGCCCTGGGATATTGCAGCCCTCCCAGACCCTCAGTCTGACTTGTTCT<br>TTCTCTGGGTTTTCACTGAGCACTTATGGTATAGGAGTAGGCTGGATTCGTCAGCCTTCAGGGAAGGG<br>TCTGGAGTGGCTGGCACACATTTGGTGGAATGATAATAAGTACTATAACACAGCCCTGAAGAGCCGG<br>CTCACAATCTCCAAGGATACCTCCAACAACCAGGTATTCCTCAAGATCGCCAGTGTGGACACTGCAGAT<br>ACTGCCACATACTACTGTGCTCGAATAG |
| IGHV-SCRF | IGHV8-13*01 | | F | CAGGTTACTCTGAAAGAGTCTGGCCCTGGTATATTGCAGCCCTCCCAGACCCTCAGTCTGACCTGTTCT<br>TTCTCTGGGTTTTCACTGAGCACTTTTGGTATGGGTGTGAGCTGGATTCGTCAGCCTTCAGGGAAGGG<br>TCTGGAGTGGCTGGCACACATTTATTGGGATGATGACAAGCACTATAACCCATCCTTGAAGAGCCGGC<br>TCACAATCTCCAAGGATACCTCCAACAACCAGGTATTCCTCAAGATCACCACTGTGGACACTGCAGATA<br>CTGCCACATACTACTGTGCTCGAAGAG |
| IGHV-SF4U | | JACKSON: balbIGHV005 | F | GAGGTCCAGCTGCAGCAGTCTGGACCTGACCTGGTGAAGCCTGGGGCTTCAGTGAAGATATCCTGCA<br>AGGCTTCTGGTTACTCATTCACTGGCTACTACATGCACTGGGTGAAGCAGAGCCATGGAAAGAGCCTT<br>GAGTGGATTGGACGTGTTAATCCTAACAATGGTGGTACTAGCTACAACCAGAAGTTCAAGGGCAAGG<br>CCATATTAACTGTAGACAAGTCATCCAGCACAGCCTACATGGAGCTCCGCAGCCTGACATCTGAGGAC<br>TCTGCGGTCTATTACTGTGCAAGA |
| IGHV-SR2I | | VBASE2: musIGHV591 | F | CAGGCTTATCTACAGCAGTCTGGGGCTGAGCTGGTGAGGTCTGGGGCCTCAGTGAAGATGTCCTGCA<br>AGGCTTCTGGCTACACATTTACCAGTTACAATATGCACTGGGTAAAGCAGACACCTGGACAGGGCCTG<br>GAATGGATTGGATATATTTATCCTGGAAATGGTGGTACTAACTACAATCAGAAGTTCAAGGGCAAGGC<br>CACATTGACTGCAGACACATCCTCCAGCACAGCCTACATGCAGATCAGCAGCCTGACATCTGAAGACT<br>CTGCGGTCTATTTCTGTGCAAGA |
| IGHV-STIW | IGHV15-2*02 | | F | CAGGTTCACCTACAACAGTCTGGTTCTGAACTGAGGAGTCCTGGGTCTTCAGTAAAGCTTTCATGCAA<br>GGATTTTGATTCAGAAGTCTTCCCTATTGCTTATATGAGTTGGGTTAGGCAGAAGCCTGGGCATGGAT<br>TTGAATGGATTGGAGACATACTCCCAAGTATTGGTAGAACAATCTATGGAGAGAAGTTTGAGGACAA |

| | | | | |
|---|---|---|---|---|
| | | | | AGCCACACTGGATGCAGACACAGTGTCCAACACAGCCTACTTGGAGCTCAACAGTCTGACATCTGAGG<br>ACTCTGCTATCTACTACTGTGCAAGG |
| IGHV-SX5D | | Haines: J558.36 | F | CCGGTCCAACTGCAGCAGCCTGGGGCTGAGCTTGTGAAGCCTGGGGCTTCAGTAAAGCTGTCCTGCA<br>AGGCTTCTGGCTACACCTTCACCAGCTACTGGATGCACTGGGTGAAGCAGAGGCCTGGACGAGGCCTT<br>GAGTGGATTGGAAGGATTGATCCTAATAGTGGTGGTACTAAGTACAATGAGAAGTTCAAGAGCAAGG<br>CCACACTGACTGTAGACAAACCCTCCAGCACAGCCTACATGCAGCTCAGCAGCCTGACATCTGAGGAC<br>TCTGCGGTCTATTACTGTACAAGA |
| IGHV-T2W5 | IGHV1S82*01 | | F | CAGGTCCAACTGCAGCAGCCTGGGGCTGAGCTGGTGAGGCCTGGAGCTTCAGTGAAGCTGTCCTGCA<br>AGGCTTCTGGCTACTCCTTCACCAGCTACTGGATGAACTGGGTGAAGCAGAGGCCTGGACAAGGCCTT<br>GAGTGGATTGGCATGATTCATCCTTCCGATAGTGAAACTAGGTTAAATCAGAAGTTCAAGGACAAGGC<br>CACATTGACTGTAGACAAATCCTCCAGCACAGCCTACATGCAACTCAGCAGCCCGACATCTGAGGACT<br>CTGCGGTCTATTACTGTGCAAGA |
| IGHV-T3DL | IGHV3-1*02 | | F | GATGTGCAGCTTCAGGAGTCAGGACCTGACCTGGTGAAACCTTCTCAGTCACTTTCACTCACCTGCACT<br>GTCACTGGCTACTCCATCACCAGTGGTTATAGCTGGCACTGGATCCGGCAGTTTCCAGGAAACAAACT<br>GGAATGGATGGGCTACATACACTACAGTGGTAGCACTAACTACAACCCATCTCTCAAAAGTCGAATCT<br>CTATCACTCGAGACACATCCAAGAACCAGTTCTTCCTGCAGTTGAATTCTGTGACTACTGAGGACACAG<br>CCACATATTACTGTGCAAGA |
| IGHV-TC2W | IGHV9-3*03 | | F | CAGATCCAGTTGGTGCAGTCTGGACCTGAGCTGAAGAAGCCTGGAGAGACAGTCAAGATCTCCTGCA<br>AGGCTTCTGGGTATACCTTCACAAACTATGGAATGAACTGGGTGAAGCAGGCTCCAGGAAAGGGTTT<br>AAAGTGGATGGGCTGGATAAACACCAACACTGGAGAGCCAACATATGCTGAAGAGTTCAAGGGACG<br>GTTTGCCTTCTCTTTGGAAACCTCTGCCAGCACTGCCTATTTGCAGATCAACAACCTCAAAAATGAGGA<br>CACGGCTACATATTTCTGTGCAAGA |
| IGHV-TI7Q | | JACKSON: balbIGHV026 | F | CAGGTTACTCTGAAAGAGTCTGGCCCTGGGATATTGCAGCCCTCCCAGACCCTCAGTCTGACTTGTTCT<br>TTCTCTGGGTTTTCACTGAGCACTTCTGGTATGGGTGTAGGCTGGATTCGTCAGCCTTCAGGGAAGGG<br>TCTGGAGTGGCTGGCACACATTTGGTGGGATGATGACAAGCGCTATAACCCAGCCCTGAAGAGCCGA<br>CTGACAATCTCCAAGGATACCTCCAGCAACCAGGTATTCCTCAAGATCGCCAGTGTGGACACTGCAGA<br>TACTGCCACATACTACTGTGCTCGAATAG |
| IGHV-TQES | | VBASE2: musIGHV672 | F | CAGGTTATTCTGAAAGAGTCTGGCCCTGGAATATTGCAGCCCTCTCAGACCCTCAGTCTGACTTGTTCT<br>TTCTCTGGGTTTTCACTTAGCACTTATGGTACAGCTGTGAACTGGATTCGTCAGCCTTCAGGAAAGGGT<br>CTGGAGTGGTTGGCACAAATTGGGTCAGATGATAGCAAGCTCTATAACCCATTTCTGAAAAGCCGAAT<br>CACAATCTCCAAGGATACCTCCAACAGCCAGGTATTCCTCAAGATCACTAGTGTGGACACTGAAGATTC<br>TGCCACATACTACTGTGCTAACAGA |

| | | | | |
|---|---|---|---|---|
| IGHV-TUT5 | | VBASE2: musIGHV585 | F | CAGGTCCAACTGCAGCAGCCTGGGGCTGAACTGGTGAAGCCTGGGGCTTCAGTGAAGCTGTCCTGCA<br>AGGCTTCTGGCTACACCTTCACCAGCTACTGGATGCACTGGGTGAAGCAGAGGCCTGGACAAGGCCTT<br>GAGTGGATTGGAGAGATTAATCCTAGCAACGGTCGTACTAACTACAATGAGAAGTTCAAGAGCAAGG<br>CCACACTGACTGTAGACAAATCCTCCAGCACAGCCTACATGCAACTCAGCAGCCTGACATCTGAGGAC<br>TCTGCGGTCTATTACTGTGCAAGA |
| IGHV-UDF7 | | JACKSON: balbIGHV039 | F | GAGGTCCAGCTGCAACAGTCTGGACCTGAGCTGGTGAAGCCTGGAGCTTCAATGAAGATATCCTGCA<br>AGGCTACTCATTCACTGGCTACACCATGAACTGGGTGAAGCAGAGCCATGGAAAGAACCTTGAGTGG<br>ATTGGACTTATTAATCCTTACAATGGTGGTACTAGCTACAACCAGAAGTTCAAGGGCAAGGCCACATT<br>AACTGTAGACAAGTCATCCAGCACAGCCTACATGGAGCTCCTCAGTCTGACATCTGAGGACTCTGCAG<br>TCTATTACTGT |
| IGHV-UJTO | IGHV1S68*02 | | F | CAGGTCCAACTGCAGCAGCCTGGGGCTGAGCTTGTGAAGCCTGGGACTTCAGTGAAGCTGTCCTGCA<br>AGGCTTCTGGCTACAACTTCACCAGCTACTGGATAAACTGGGTGAAGCTGAGGCCTGGACAAGGCCTT<br>GAGTGGATTGGAGATATTTATCCTGGTAGTGGTAGTACTAACTACAATGAGAAGTTCAAGAGCAAGG<br>CCACACTGACTGTAGACACATCCTCCAGCACAGCCTACATGCAACTCAGCAGCCTGGCATCTGAGGAC<br>TCTGCTCTCTATTACTGTGCAAGA |
| IGHV-UOVB | IGHV14-3*02 | | F | GAGGTTCAGCTGCAGCAGTCTGGGGCAGAGCTTGTGAAGCCAGGGGCCTCAGTCAAGTTGTCCTGCA<br>CAGCTTCTGGCTTCAACATTAAAGACACCTATATGCACTGGGTGAAGCAGAGGCCTGAACAGGGCCTG<br>GAGTGGATTGGAAGGATTGATCCTGCGAATGGTAATACTAAATATGACCCGAAGTTCCAGGGCAAGG<br>CCACTATAACAGCAGACACATCCTCCAACACAGCCTACCTGCAGCTCAGCAGCCTGACATCTGAGGAC<br>ACTGCCGTCTATTACTGTGCTAGA |
| IGHV-UTQI | IGHV5-6-4*01 | | F | GACGTGAAGCTGGTGGAGTCTGGGGGAGGCTTAGTGAAGCCTGGAGGGTCCCTGAAACTCTCCTGTG<br>CAGCCTCTGGATTCACTTTCAGTAGCTATACCATGTCTTGGGTTCGCCAGACTCCGGAGAAGAGGCTG<br>GAGTGGGTCGCAACCATTAGTAGTGGTGGTAGTTACACCTACTATCCAGACAGTGTGAAGGGCCGATT<br>CACCATCTCCAGAGACAATGCCAAGAACACCCTGTACCTGCAAATGAGCAGTCTGAAGTCTGAGGACA<br>CAGCCATGTATTACTGTACAAGAGA |
| IGHV-V26Z | IGHV2-9*02 | | F | CAGGTGCAGCTGAAGGAGTCAGGACCTGGCCTGGTGGCGCCCTCACAGAGCCTGTCCATCACTTGCA<br>CTGTCTCTGGGTTTTCATTAACCAGCTATGGTGTACACTGGGTTCGCCAGCCTCCAGGAAAGGGTCTG<br>GAGTGGCTGGGAGTAATATGGGCTGGTGGAAGCACAAATTATAATTCGGCTCTCATGTCCAGACTGA<br>GCATCAGCAAAGACAACTCCAAGAGCCAAGTTTTCTTAAAAATGAACAGTCTGCAAACTGATGACACA<br>GCCATGTACTACTGTGCCAGAGA |
| IGHV-W27J | | VBASE2: musIGHV555 | F | CACGTCCAGCTGCAGCAATCTGGACCTGAGCTGGTGAGGCCTGGGGCTTCAGTGAAGCTGTCCTGCA<br>AGGCTTCTGGCTATATCTTCATCACCTACTGGATGAACTGGGTGAAGCAGAGGCCTGGACAGGGCCTT<br>GAGTGGATTGGACAGATTTTTCCTGCAAGTGGTAGTACTAACTACAATGAGATGTTCGAGGGCAAGG |

| | | | | |
|---|---|---|---|---|
| | | | | CCACATTGACTGTAGACACATCCTCCAGCACAGCCTACATGCAGCTCAGCAGCCTGACATCTGAGGAC<br>TCTGCGGTCTATTACTGTGCAAGA |
| IGHV-WN4M | IGHV2-6-7*01 | | F | CAGGTGCAGCTGAAGGAGTCAGGACCTGGCCTGGTGGCGCCCTCACAGAGCCTGTCCATCACATGCA<br>CCGTCTCAGGGTTCTCATTAACCGGCTATGGTGTAAACTGGGTTCGCCAGCCTCCAGGAAAGGGTCTG<br>GAGTGGCTGGGAATGATATGGGGTGATGGAAGCACAGACTATAATTCAGCTCTCAAATCCAGACTGA<br>GCATCAGCAAGGACAACTCCAAGAGCCAAGTTTTCTTAAAAATGAACAGTCTGCAAACTGATGACACA<br>GCCAGGTACTACTGTGCCAGAGA |
| IGHV-WO3N | | JACKSON: balbIGHV029 | F | CAGGTCCAACTGCAGCAGTCTGGGGCTGAACTGGTGAAGCCTGGGGCTTCAGTGAAGTTGTCCTGCA<br>AGGCTTCTGGCTACACCTTCACCAGCTACTATATGTACTGGGTGAAGCAGAGGCCTGGACAAGGCCTT<br>GAGTGGATTGGAGAGATTAATCCTAGCAATGGTGGTACTAACTTCAATGAGAAGTTCAAGAGCAAGG<br>CCACACTGACTGTAGACAAATCCTCCAGCACAGCATACATGCAACTCAGCAGCCTGACATCTGAGGAC<br>TCTGCGGTCTATTACTGTACAAGA |
| IGHV-WZ7P | IGHV2-6-2*01 | | F | CAGGTGCAGCTGAAGGAGTCAGGACCTGACCTGGTGGCGCCCTCACAGAGCCTGTCCATCACATGCA<br>CCGTCTCAGGGTTCTCATTAACCAGCTATGGTGTACACTGGGTTCGCCAGCCTCCAGGAAAGGGTCTG<br>GAGTGGCTGGTAGTGATATGGAGTGATGGAAGCACAACCTATAATTCAGCTCTCAAATCCAGACTGA<br>GCATCAGCAAGGACAACTCCAAGAGCCAAGTTTTCTTAAAAATGAACAGTCTCCAAACTGATGACACA<br>GCCATGTACTACTGTGCCAGACA |
| IGHV-X4X2 | | Haines: J558.27 | F | CAGGTCCAGCTGCAGCAGTCTGGACCTGAGCTGGTGAAGCCTGGGGCTTCAGTGAAGATGTCCTGCA<br>AGGCTTCTGGCTACACCTTCACAAGCTACTATATACACTGGGTGAAGCAGAGGCCTGGACAGGGACTT<br>GAGTGGATTGGATGGATTTATCCTGGAGATGGTAGTACTAAGTACAATGAGAAGTTCAAGGGCAAGA<br>CCACACTGACTGCAGACAAATCCTCCAGCACAGCCTACATGTTGCTCAGCAGCCTGACCTCTGAGGACT<br>CTGCGATCTATTTCTGTGCAAGA |
| IGHV-XGYI | IGHV5-12-1*01 | | F | GAAGTGCAGCTGGTGGAGTCTGGGGGAGGCTTAGTGAAGCCTGGAGGGTCCCTGAAACTCTCCTGTG<br>CAGCCTCTGGATTCGCTTTCAGTAGCTATGACATGTCTTGGGTTCGCCAGACTCCGGAGAAGAGGCTG<br>GAGTGGGTCGCATACATTAGTAGTGGTGGTGGTAGCACCTACTATCCAGACACTGTGAAGGGCCGAT<br>TCACCATCTCCAGAGACAATGCCAAGAACACCCTGTACCTGCAAATGAGCAGTCTGAAGTCTGAGGAC<br>ACAGCCATGTATTACTGTGCAAGACA |
| IGHV-YEPS | | JACKSON: balbIGHV036 | F | CAGGTTACTCTGAAAGAGTCTGGCCCTGGGATATTGCAGCCATCACAGACGCTTAGCCTGGCCTGTAC<br>TTTCTCTGGGATTTCACTGAGTACTTCTGGTATGGGTTTGAGCTGGCTTCGTAAGCCCTCAGGGAAGGC<br>TTTAGAGTGGCTGGCAAGCATTTGGAATAATGATAACTACTACAACCCATCTTTGAAGAGCCGGCTCA<br>CAATCTCCAAGGAGACCTCCAACAACCAAGTATTCCTTAAACTCACCAGTGTGGACACTGCAGATTCTG<br>CCACATACTACTGTGCTTGGAGAGAG |

| | | | | |
|---|---|---|---|---|
| IGHV-Z6OV | IGHV1S72*01 | | F | CAGGTCCAACTGCAGCAGCCTGGGGCTGAGCTTGTGAAGCCTGGGGCTCCAGTGAAGCTGTCCTGCA AGGCTTCTGGCTACACCTTCACCAGCTACTGGATGAACTGGGTGAAGCAGAGGCCTGGACGAGGCCT CGAGTGGATTGGAAGGATTGATCCTTCCGATAGTGAAACTCACTACAATCAAAAGTTCAAGGACAAGG CCACACTGACTGTAGACAAATCCTCCAGCACAGCCTACATCCAACTCAGCAGCCTGACATCTGAGGACT CTGCGGTCTATTACTGTGCAAGA |
| IGHV-ZD3D | IGHV2-5-1*01 | | F | CAGGTGCAGCTGAAGCAGTCAGGACCTAGCCTAGTGCAGCCCTCACAGAGCCTGTCCATAACCTGCAC AGTCTCTGGTTTCTCATTAACTAGCTATGGTGTACACTGGGTTCGCCAGTCTCCAGGAAAGGGTCTGGA GTGGCTGGGAGTGATATGGAGAGGTGGAAGCACAGACTACAATGCAGCTTTCATGTCCAGACTGAGC ATCACCAAGGACAACTCCAAGAGCCAAGTTTTCTTTAAAATGAACAGTCTGCAAGCTGATGACACTGC CATATACTACTGTGCCAAAA |
| IGHV-ZGQJ | IGHV5-9-2*01 | | F | GAAGTGAAGCTGGTGGAGTCTGGGGGAGGCTTAGTGAAGCCTGGAGGGTCCCTGAAACTCTCCTGTG CAGCCTCTGGATTCACTTTCAGTAGCTATGGCATGTCTTGGGTTCGCCAGACTCCGGAGAAGAGGCTG GAGTGGGTCGCAACCATTAGTGGTGGTGGTAGTTACACCTACTATCCAGACAGTGTGAAGGGGCGAT TCACCATCTCCAGAGACAATGCCAAGAACAACCTGTACCTGCAAATGAGCAGTCTGAGGTCTGAGGAC ACGGCCTTGTATTACTGTGCAAGACA |
| IGHV-ZI2X | IGHV2-4*02 | | F | CAGGTGCAGCTGAAGCAGTCAGGACCTGGCCTAGTGCAGCCCTCACAGAGCCTGTCCATCACCTGCAC AGTCTCTGGTTTCTCATTAACTAGCTATGGTGTACACTGGGTTCGCCAGCCTCCAGGAAAGGGTCTGG AGTGGCTGGGAGTGATATGGAGTGGTGGAAGCACAGACTATAATGCTGCTTTCATATCCAGACTGAG CATCAGCAAGGACAACTCCAAGAGCCAAGTTTTCTTTAAAATGAACAGTCTGCAAGCTGATGACACAG CCATATACTACTGTGCCAGAAA |
| IGHV-ZIT7 | | VBASE2: musIGHV655 | F | CAGGTCCAACTGCAGCAGCCTGGGGCTGAGCTGGTGAGGCCTGGGGCTTCAGTGAAGCTGTCCTGCA AGGCTTCTGGCTACACCTTCACCAGCTACTGGATAAACTGGGTGAAGCAGAGGCCTGGACAAGGCCTT GAGTGGATCGGAAATATTTATCCTTCTGATAGTTATACTAACTACAATCAAAAGTTCAAGGACAAGGC CACATTGACTGTAGACAAATCCTCCAGCACAGCCTACATGCAGCTCAGCAGCCCGACATCTGAGGACT CTGCGGTCTATTACTGTACAAGA |
| IGHV-ZL77 | IGHV3-5*02 | | F | GATGTGCAGCTTCAGGAGTCAGGACCTGGTCTGGTGAAACCTTCTCAGACAGTGTCCCTCACCTGCAC TGTCACTGGCATCTCCATCACCACTGGAAATTACAGATGGAGCTGGATCCGGCAGTTTCCAGGAAACA AACTGGAGTGGATAGGGTACATATACTACAGTGGTACCATTACCTACAATCCATCTCTCACAAGTCGA ACCACCATCACTAGAGACACTTCCAAGAACCAATTCTTCCTGGAAATGAACTCTTTGACTGCTGAAGAC ACAGCCACATACTACTGTGCACGAGA |
| IGHV-ZOKF | | JACKSON: balbIGHV019 | F | GAGATCCAGCTGCAGCAGTCTGGACCTGAGCTGGTGAAGCCTGGGGCTTCAGTGAAGGTATCCTGCA AGGCTTCTGGTTATGCATTCACTAGCTACAACATGTACTGGGTGAAGCAGAGCCATGGAAAGAGCCTT GAGTGGATTGGATATATTGATCCTTACAATGGTGGTACTAGCTACAACCAGAAGTTCAAGGGCAAGGC |

| | | | |
|---|---|---|---|
| | | | CACATTGACTGTTGACAAGTCCTCCAGCACAGCCTACATGCATCTCAACAGCCTGACATCTGAGGACTCTGCAGTCTATTACTGTGCAAGA |
| IGHV-ZOZI | VBASE2: musIGHV528 | F | GAGGTCCAGCTGCAACAGTCTGGACCTGAGCTGGTGAAGCCTGGGGCTTCAGTGAAGATGTCCTGCAAGGCTTCTGGATACACCTTCACTGACTACTACATGAAGTGGGTGAAGCAGAGCCATGGAAAGAGCCTTGAGTGGATTGGAGATATTAATCCTAACAATGGTGATACTTTCTACAACCAGAAGTTCAAGGGCAAGGCCACATTGACTGTAGACAAATCCTCCAGCACAGCCTACATGCAGCTCAACAGCCTGACATCTGAGGACTCTGCAGTCTATTACTGTGCAAGA |
| IGHV-ZVAP | JACKSON: balbIGHV032 | F | CAGGTCCAGCTGCAGCAGTCTGGGCCTGAGCTGGTGAGGCCTGGGGTCTCAGTGAAGATTTCCTGCAAGGGTTCCGGCTACACATTCACTGATTATGCTATGCACTGGGTGAAGCAGAGTCATGCAAAGAGTCTAGAGTGGATTGGAGTTATTAGTACTTACTCTGGTAATACAAACTACAACCAGAAGTTTAAGGGCAAGGCCACAATGACTGTAGACAAATCCTCCAGCACAGCCTATATGGAACTTGCCAGATTGACATCTGAGGATTCTGCCATCTATTACTGTGCAAGA |
